# Assembly of two functionally-distinct protein import complexes in the outer membrane of plant chloroplasts

**DOI:** 10.1101/2025.10.27.683105

**Authors:** Sreedhar Nellaepalli, Domagoj Baretic, Astrid F. Brandner, Sybille Kubis-Waller, Ziad Soufi, Shuyang Cheng, Sireesha Kodru, Jun Fang, Ursula Flores-Perez, Vaishnavi Ravikumar, Pablo Pulido, Marjorie Fournier, Ivan Ahel, Syma Khalid, R. Paul Jarvis

## Abstract

The TOC translocon delivers thousands of nucleus-encoded proteins to chloroplasts and related non-photosynthetic plastids. It comprises the β-barrel channel, Toc75, and multiple isoforms of receptor GTPases, Toc33 and Toc159. However, exactly how TOC complexes are assembled in different plastid types is unknown. Here, we present detailed characterization of two distinct TOC complexes, TOC-P and TOC-N, from photosynthetic chloroplasts and non-photosynthetic plastids, respectively. The assembled complexes are distinguished by having different sets of receptors, but both possess Toc75 which we identify as a central hub in TOC biogenesis: assembly is driven by *TOC75* expression, with the Toc33 and Toc159 being added sequentially thereafter. Integrative structural analysis revealed a modular architecture for TOC-P comprising a cytosolic GTPase receptor module linked flexibly to a membrane β-barrel channel module. TOC-N has a similar overall architecture, albeit with some clear differences that likely account for observed functional differences related to client specificity.

## MAIN

Chloroplasts are membrane-bound endosymbiotic organelles in plants and algae that perform photosynthesis, the energetic basis for essentially all life on Earth^1^. They belong to a broader family of interconvertible organelles called the plastids, along with several non-photosynthetic variants (e.g., etioplasts in dark-grown plants, and leucoplasts in roots)^2^. Chloroplasts and other plastids are built-up from thousands of different proteins, and because these are mostly nucleus-encoded, organelle biogenesis requires the efficient operation of sophisticated protein-import machines^3–6^. This machinery enables the massive, rapid delivery of thousands of different proteins to developing chloroplasts during photosynthetic establishment.

In plants, the plastid protein import machinery comprises two multiprotein complexes called TOC and TIC (translocon of the <u>o</u>uter/inner <u>c</u>hloroplast membrane). TOC recognizes and imports preproteins from the cytosol to the intermembrane space (IMS), while TIC completes delivery to the stroma in coordination with import-motor systems^3–6^. Because TOC governs which proteins enter the chloroplast, it is the crucial gatekeeper of the import machinery. The TOC comprises a channel-protein, Toc75, and two preprotein-receptors, Toc33 and Toc159. As an Omp85-superfamily member (related to proteins in bacteria and mitochondria), Toc75 comprises a β-barrel domain that forms a preprotein-conducting channel, and a polypeptide-transport-associated (POTRA) domain that chaperones preproteins through the IMS^7^. The receptors possess related GTPase domains facing the cytosol^3^. Toc33 is structurally-simple with a single α-helical membrane-span at its C-terminus; while Toc159 is more complex with an N-terminal acidic (A) domain and a large C-terminal membrane domain either side of the central GTPase domain.

In *Arabidopsis thaliana* and other plants, the receptors exist in multiple isoforms with different properties, forming two families (i.e., Toc159, Toc132, Toc120; and Toc33, Toc34)^3,8–10^. Toc159 and Toc33 are preferentially associated with photosynthetic development in green tissues, whereas the other isoforms, Toc132, Toc120 and Toc34, are more prominent in non-photosynthetic tissues, such as roots, and may help to maintain basic organelle functions^1^. Accordingly, plant knockout (ko) mutants of Toc33 (*plastid protein import 1*, *ppi1*) and Toc159 (*ppi2*) exhibit especially pronounced effects on chloroplast biogenesis^11–15^. Existence of these different receptor types may enable nuanced regulation of protein import to dynamically control the organelle’s proteome and functions, potentially circumventing damaging competition effects among the precursors. It may also facilitate to the differentiation of photosynthetic and non-photosynthetic plastid types.

Although the composition of the TOC apparatus is generally well established, the structural characteristics of the complexes and their differences between plastid types, as well as the molecular details of the relevant assembly processes, are poorly understood^16,17^. Here we have addressed these knowledge gaps by studying different TOC configurations present in chloroplasts and non-green plastids. Using a combination of structural analysis following affinity purification and modelling, together with cross-linking, genetic, and in vivo labelling approaches, we elucidate the structural organization and functions of the two distinct spatially enriched TOC complexes, and identify Toc75 as a central hub or “master player” in the biogenesis of TOC complexes.

## RESULTS

### Isolation of two distinct TOC complexes from different plastid types

To enable analysis of TOC composition in different plastid types, we generated *A. thaliana* transgenic plants expressing Toc75 with an HA epitope tag inserted close to the N-terminus of the mature protein, between amino acid residues E142 and E143 of the precursor (E142-HA-E143); this was done in a *toc75* knockdown (kd) mutant background (i.e., *mar1/toc75-III-3*, bearing a G658R missense mutation) which accumulates Toc75 to just ∼30% of the normal level^18^ **(Fig. 1a)**. Three independent *HA-Toc75* transgenic lines showed recovery to the wild-type green phenotype **(Fig. 1b,c)**, linked to restored Toc75 protein accumulation **(Fig. 1d)**. This indicated that insertion of the HA-tag at E142, which is close to the N-terminal linker (148-172) that caps the POTRA1 domain^7^, did not affect the function of Toc75.

**Fig. 1.**
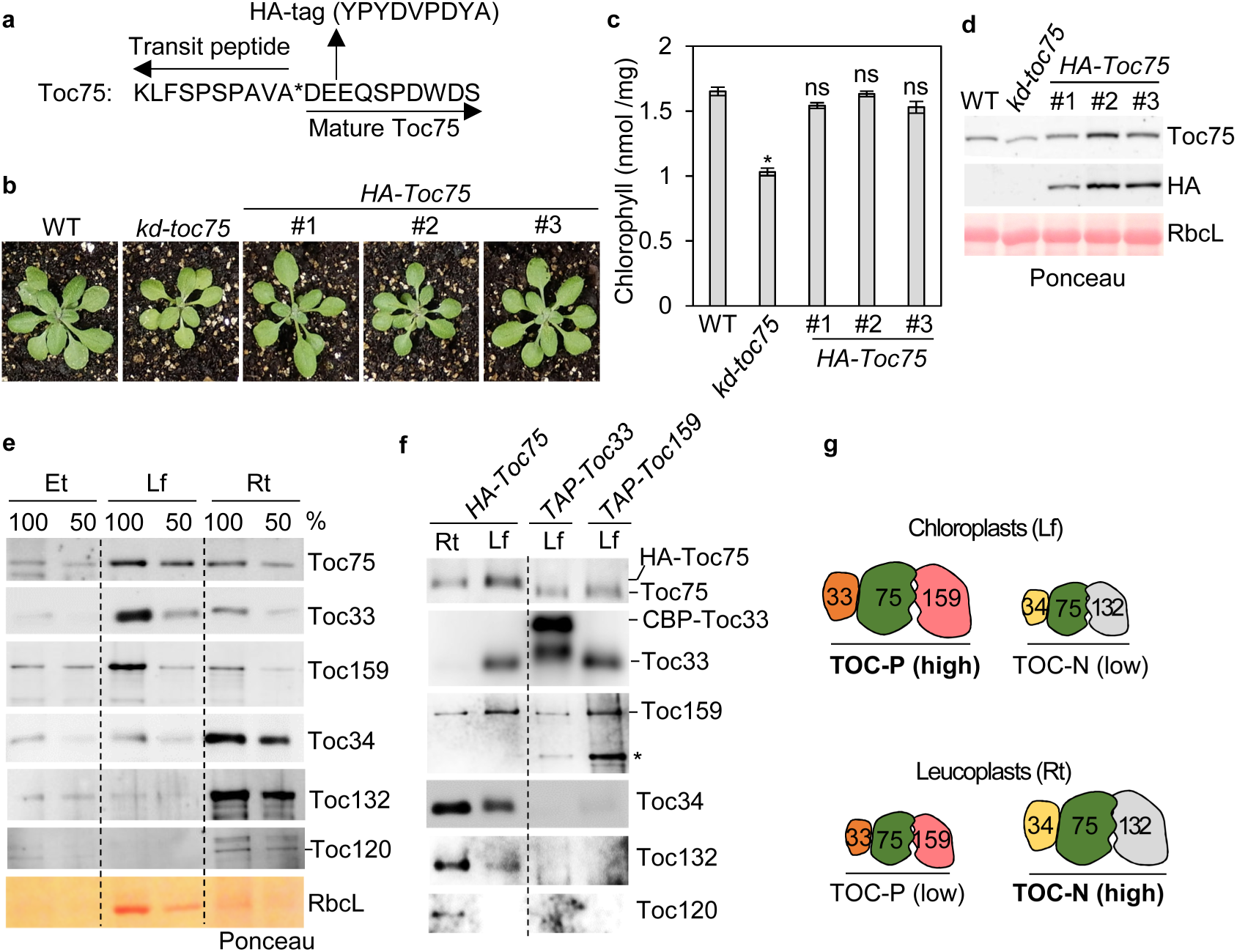
Isolation of two distinct TOC complexes, TOC-P and TOC-N. **a.** Insertion site of the HA-tag in Toc75 of the *HA-Toc75* transgenic lines (142E-HA-143E); D141 is the N-terminus of mature Toc75 after transit peptide (TP) cleavage. Asterisk indicates the TP cleavage site. **b.** Visible phenotypes of WT, *kd-toc75* and three independent *HA-Toc75* transgenic lines (*#1-3*). **c.** Chlorophyll concentrations in 4-week-old plants of the genotypes shown in **b**. Asterisks indicate significance according to paired two-tailed Student’s t tests comparing the mutant genotypes with WT (\**p* < 0.001; ns, not significant). All values are means ± SEM (n = 4 experiments). **d.** Immunoblotting analysis of total protein extracts from 12-14-day-old plants of the genotypes shown in **b**. Samples were loaded based on equal plant fresh weight. **e.** Immunoblotting analysis of protein extracts from dark-grown wild-type seedlings containing etioplasts (Et); from chloroplast-enriched leaf fractions (Lf); and from roots containing leucoplasts (Rt). The samples were prepared from 7-day-old (Et) or 14-day-old (Lf, Rt) plants. **f.** Isolation of TOC complexes by HA- or TAP-tag affinity purification from Lf or Rt samples (similar to those in **e**) from the indicated transgenic lines. Samples were solubilized with 1% β-DM before affinity purification. Purified samples were analysed by immunoblotting, with loading normalized to achieve comparable Toc75 levels. *HA-Toc75* line #1 was used for the purifications. Asterisk indicates proteolytic fragment of Toc159. **g.** Schematic representation of the TOC-P and TOC-N complexes that are enriched in chloroplasts and leucoplasts, respectively. Although TOC-P and TOC-N are also present in leucoplasts and chloroplasts, respectively, the abundance is low compared to the major complex in each plastid type.

Initially, we determined the abundance of TOC components in different tissues of the wild type by immunoblotting. Receptor isoforms Toc33 and Toc159 were enriched in leaves (in chloroplasts), whereas Toc34 and Toc132/120 (Toc132 and Toc120 are highly similar and considered to be largely or completely redundant^15^) were enriched in roots and petals (in non-green plastids or leucoplasts), revealing clear differentiation of TOC composition in different plastid types **(Fig. 1e and Supplementary Fig. 1)**. Interestingly, all TOC components were present at relatively low levels in etiolated seedlings, which contain partially differentiated plastids called etioplasts.

Next, we applied anti-HA affinity chromatography to purify TOC complexes from different tissues of the *HA-Toc75* transgenic plants^19^. This analysis showed that Toc75 is mainly associated with Toc33 and Toc159 in leaf chloroplasts, and to a lesser extent with Toc34, Toc132 and Toc120 **(Fig. 1f)**. In contrast, Toc75 is mainly associated with Toc34, Toc132 and Toc120 in root leucoplasts, and to a lesser extent with Toc33 and Toc159 **(Fig. 1f, left side)**. These data are consistent with the aforementioned TOC accumulation biases **(Fig. 1e)**, and point to the predominance of distinct TOC complexes in leaf chloroplasts and root leucoplasts, although clearly the prevalence of one type does not preclude co-existence of the other at low levels. For clarity, we herein refer to the Toc33-75-159 configuration as TOC-P (for photosynthetic) and the Toc34-75-132/120 configuration as TOC-N (for non-photosynthetic) **(Fig. 1g)**.

We wished to determine if TOC-P and TOC-N interact in chloroplast membranes. For this purpose, we used transgenic lines expressing tandem-affinity-purification (TAP)-tagged Toc33 and Toc159 (*TAP-Toc33* and *TAP-Toc159*)^20,21^ to affinity purify TAP-Toc33 and TAP-159 complexes, respectively, from chloroplast extracts **(Fig. 1f, right side)**. In the resulting samples, co-purification of Toc34 and Toc132 was negligible by comparison with Toc33 and Toc159; whereas in affinity-purified HA-Toc75 complexes, Toc34 and Toc132 were both clearly detectable **(Fig. 1f, left side)**. This indicated that TOC-P and TOC-N exist as distinct complexes and that Toc75 is the common factor of both complexes.

### Toc75 and Toc33/34 maintain the stability of TOC complexes

To elucidate which TOC components are important in stabilizing the integrity of the TOC complexes, we analysed the accumulation of all TOC components in the following *Arabidopsis* TOC mutant genotypes: *mar1/toc75-III-3* (*kd-toc75*)^22,23^, *ppi1-1* (*ko*-*toc33*)^14^, *ppi3-2* (*ko-toc34*)^12^, *fts1/ppi2-3* (*kd-toc159*)^20,24^, and *toc132-2* (*ko-toc132*)^15^ **(Fig. 2)**. As previously described, these mutants typically display a characteristic chlorotic phenotype **(Supplementary Fig. 2)**.

**Fig. 2.**
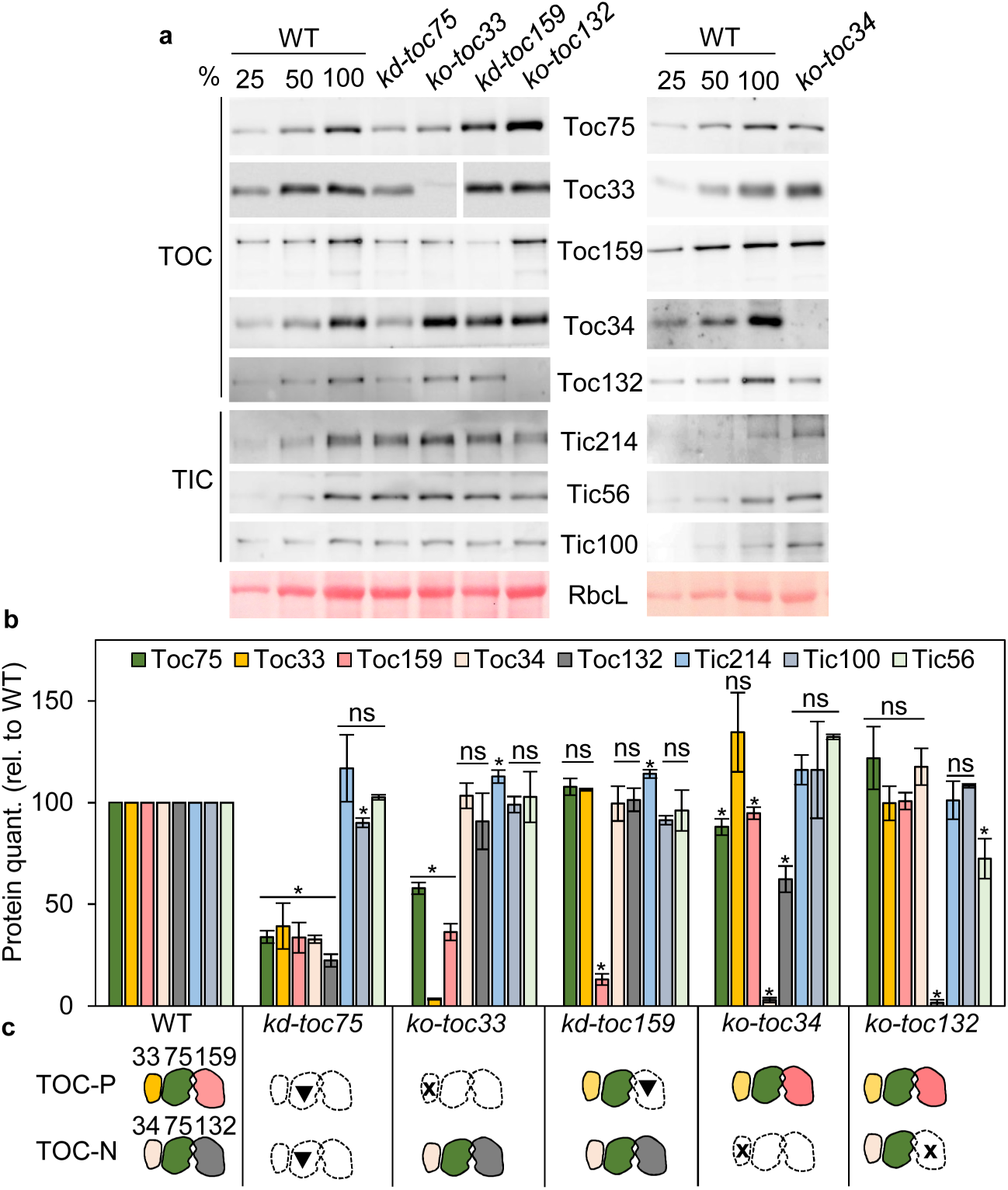
Comparison of the contributions of different TOC components to TOC complex stability. **a.** Immunoblotting analysis of leaf protein extracts from 12-14-day-old WT, *kd-toc75* (*mar1/toc75-III-3*), *ko-toc33* (*ppi1-1*), *kd-toc159* (*fts1/ppi2-3*), *ko-toc132* (*toc132-2*), and *ko-toc34* (*ppi3-2*) plants. Samples were loaded based on equal plant fresh weight. **b.** Protein quantification of the immunoblots in a and of other similar results. All values are means ± SEM (n = 3-4 experiments). Asterisks indicate significance according to paired one-tailed Student’s t tests comparing the mutant genotypes with WT (\**p* < 0.02; ns, not significant). **c.** Schematic representation of the results in **a** and **b**. Dotted lines indicate reduced accumulation of the relevant component. Components marked x are affected by a knockout (ko) mutation; components marked with a triangle are affected by a knockdown (kd) mutation.

As reported previously^18^ and shown earlier **(Fig. 1b)**, the *kd-toc75* mutant displayed a ∼70% reduction in Toc75 accumulation **(Fig. 2a,b)**. In this mutant, the Toc33/Toc34 and Toc159/Toc132 receptor isoforms were all similarly reduced, revealing strong effects on the integrity of both TOC-P and TOC-N. In contrast, in the *ko-toc33* mutant, Toc75 and Toc159 were both reduced by ∼50% but the components of TOC-N (Toc34 and Toc132) were unaffected. This implies that TOC-P is specifically disrupted in *ko-toc33*. Similarly, in the *ko-toc34* mutant, Toc132 was the most strongly affected protein amongst the other TOC components, indicating a specific effect on TOC-N. Interestingly, in the *kd-toc159* and *ko-toc132* mutants, the Toc75, Toc33, and Toc34 proteins were not affected, indicating that Toc159 and Toc132 are not required for the stability of the other components of TOC-P and TOC-N, respectively. In general, TIC components were largely unaffected in the different *toc* mutants **(Fig. 2a,b)**. Thus, in summary, these results indicate that Toc75 and Toc33/34 are the key players in stabilizing the integrity of the TOC-P and TOC-N complexes **(Fig. 2c)**.

Importantly, qRT-PCR analysis revealed that *TOC* gene expression changes in the *toc* mutants were minimal, and generally positive (not negative) **(Supplementary Fig. 3)**. This was the case even in the *kd-toc75* and *ko-toc33* mutants which displayed the strongest effects on TOC protein accumulation. This supports the view that the observed protein effects **(Fig. 2c)** were due to defects at the post-translational level.

### Integrative structural analysis reveals a bipartite structure for TOC-P

To gain insight into the structural basis for our biochemical observations **(Figs. 1 and 2)**, we implemented an integrative structural analysis. First, a computational approach using AlphaFold3 (AF3) was applied to analyse the TOC-P complex^25^. The amino acid sequences of Toc75 (without the transit peptide), Toc33, and Toc159 (without the unstructured A domain^26^) from *A. thaliana*, plus two GTP ligands, were submitted to the AF3 server to generate structural predictions **(Fig. 3a)**; the sequence of the Toc159 A domain was removed as its intrinsically disordered nature poses a challenge for AF3 prediction^26^. We then analysed the top-scoring TOC model to elucidate domain organization and component interactions within the complex.

**Fig. 3.**
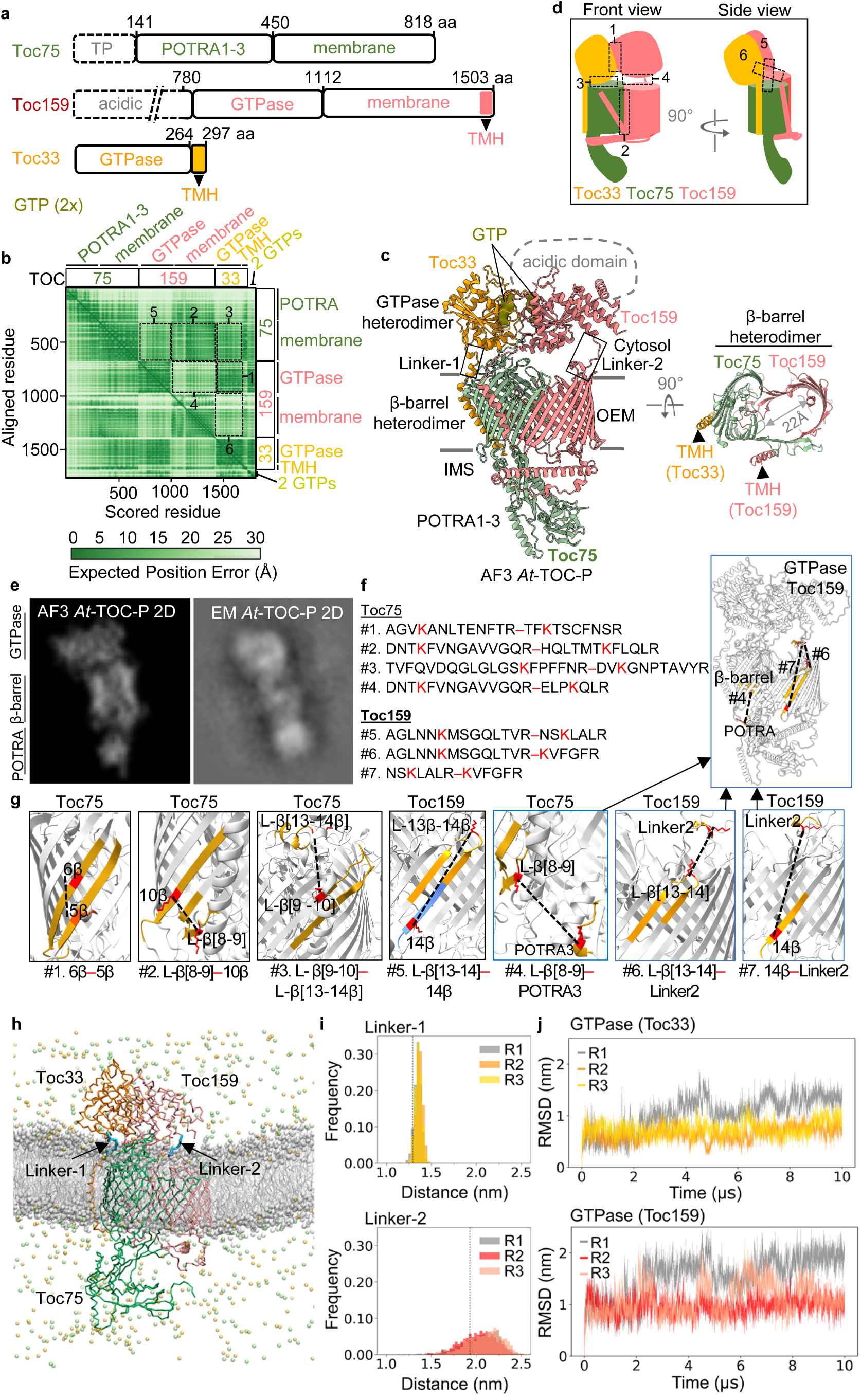
Integrative structural analysis of TOC-P reveals a heterodimeric-GTPase–heterodimeric-β-barrel architecture. **a.** Components and ligands (two GTPs) of the TOC-P complex submitted to the AF3 server. Domain maps of the inputted protein sequences are shown; the sequences of Toc75 and Toc159 were N-terminally truncated to remove transit peptide and disordered acidic A-domain sequences, respectively. The numbers refer to amino acid (aa) coordinates of the full-length proteins. **b.** Predicted Alignment Error (PAE) plot from the AF3 analysis of the components in **a**. This 2D plot shows expected position error (Å) of different domains and interfaces, and measures the expected positional error (confidence) of pairs of positions running along the axes. Dark green indicates lower error, while light green indicates higher error. Dashed boxes 1-6 indicate interfaces between the TOC subunits as follows: 1, Toc33-GTPase:Toc159-GTPase; 2, Toc75-membrane:Toc159-membrane; 3, Toc33-GTPase:Toc75-membrane; 4, Toc159-GTPase:Toc159-membrane; 5, Toc75-membrane:Toc159-GTPase; 6, Toc159-membrane:Toc33-GTPase. These interfaces are also marked in **d**. **c.** Structure of the TOC-P complex as modelled by AF3 (front and top views). In the front view (left), the position of the acidic domain of Toc159 (which was excluded from the analysis because of its intrinsically disordered nature) is shown by the dashed line. Two linkers connecting the GTPase heterodimer with the β-barrel heterodimer are identified using rectangles. In the top view (90° rotation relative to front view), the membrane domain of TOC-P, composed of a Toc75-Toc159 hybrid channel, is clearly shown. Transmembrane helices (TMH) of Toc33 and Toc159 are also shown. **d.** Simplified front and side view representations of the TOC-P structure shown in **c**. **e.** Comparison of a 2D back projection of the AF3-derived TOC-P structure shown in **c** (generated by CryoSPARC) (left) with a similar 2D class average from the negative staining EM analysis of purified TOC-P complex (right). **f.** XL-MS analysis of the TOC-P complex identified crosslinks in Toc75 (#1-4) and Toc159 (#5-7). The affinity purified complex was crosslinked with BS3 and trypsin digested prior to MS analysis. Crosslinked peptides are shown, with the crosslink highlighted in red in each case. For full list of identified peptides, refer to **Supplementary Table 1**. **g.** Each crosslink detailed in **f** (#1-7) was mapped onto the TOC-P structure from **c**. In each image, crosslinked parts of the structure are highlighted (yellow for peptides, red for residues; for #5, one peptide is in blue as the two peptides are contiguous), with the crosslinks shown by the dotted black lines. Positions of interdomain crosslinks (marked with blue boxes) are shown in the larger image at right. Linker-2 connects the GTPase and β-barrel domains of Toc159. L, loop. **h.** Setup for the molecular dynamics (MD) simulation analysis of TOC-P. The complex from c was embedded in a symmetric 1-palmitoyl-2-oleoyl-*sn*-glycero-3-phosphocholine (POPC) lipid bilayer. The protein chains are coloured as in c; POPC molecules are shown in light grey with their headgroup moieties (phosphate and choline groups) represented as spheres; and Cl^−^ and Na^+^ ions are shown as green and orange spheres, respectively. Water is omitted for clarity. Three independent simulation runs (R1-R3) were performed. **i.** Flexibility of linker-1 and linker-2 in the MD analysis. Histograms showing the end-to-end distance distribution during the full production run for all three independent replicates. The dotted vertical line in each case corresponds to the value of the initial AF3 model. **j.** Mobility of the GTPase domains of Toc33 and Toc159 in the MD analysis. RMSD plots of the GTPase domains after fitting on the transmembrane region of the TOC-P complex, for all three independent runs.

AF3 predicted the organization of the TOC-P complex with high confidence (ipTM = 0.72; pTM = 0.74) **(Fig. 3b)**. The model revealed a striking heterodimeric-GTPase–heterodimeric-β-barrel structure, as follows: the cytosolic GTPase domains of Toc33 and Toc159 form a heterodimeric receptor module connected by two linkers to an integrated membrane channel module, the latter composed primarily of the β-barrel segments of Toc75 and Toc159 which together form a heterodimeric hybrid barrel; Toc75 further extends a globular domain comprising three POTRA domains into the presumptive IMS. Overall, TOC-P forms a bipartite receptor-channel complex with a novel arrangement **(Fig. 3c,d)**.

To validate the presented TOC-P model, we performed negative-staining analysis of purified TOC-P complex by electron microscopy (EM) **(Fig. 3e)** and crosslinking mass-spectrometry (XL-MS) **(Fig. 3f,g)**. For the former, TOC-P complex was purified from chloroplast membrane extracts of the *HA-Toc75* transgenic line **(Fig. 1)** by affinity purification and size-exclusive chromatography, and then negatively stained prior to EM imaging and 2D classification of the images. The results revealed that purified TOC-P single particles have a very similar modular arrangement to that seen in the TOC-P AF3 model, as revealed by comparison with 2D projections of the AF3 model **(Fig. 3e and Extended Data Fig. 1)**. For the XL-MS analysis, a similar affinity-purified TOC-P sample was chemically crosslinked using BS3 prior to proteolytic digestion for MS analysis. This identified several crosslinks within Toc75 and Toc159 **(Fig. 3f)**, and the positioning of each of these crosslinks was accordant with AF3 model as the relevant residues were all in close proximity within the structure **(Fig. 3g)**. Notably, crosslinks between the POTRA and β-barrel domains of Toc75 (crosslink #4), and between the central linker and β-barrel domain of Toc159 (crosslinks #6 and #7), show close proximity between these elements of the structure **(Fig. 3g)**. In summary, these EM and XL-MS results support the veracity of the TOC-P AF3 model. Such an integrative approach, combining computational modelling with experimental validation as described, may present significant advantages over solely cryo-EM-focused approaches when analysing complexes with flexible domains such as the TOC complex (see below)^27,28^.

### AF3 analysis reveals a similar structure for TOC-N

Next, we performed a similar analysis of the TOC-N complex by AF3 which also resulted in a high-confidence model (ipTM = 0.70; pTM = 0.72) **(Extended Data Fig. 2a,b)**. As might be expected given its functional similarity, TOC-N shares similar structural features with TOC-P (i.e., a heterodimeric-GTPase–heterodimeric-β-barrel arrangement). However, the two complexes do exhibit some notable differences, including an extended IMS helix in Toc34 **(Extended Data Fig. 2b)**, a disordered α5’ region in Toc34 **(Extended Data Fig. 2c)**, and several areas of contrasting surface electrostatic potential **(Extended Data Fig. 2f)**. Such differences could be partly responsible for functional differentiation between TOC-P and TOC-N, for example in relation to possible client specificity differences. Although the A-domains of Toc159 (aa 1-786) and Toc132 (aa 1-485) were not analyzed here, because their intrinsically disordered nature presents problems for structural analysis, these domains may also be important in defining client specificities^13,26^.

### Details of the TOC-P model reveal modular assembly and intimate Toc75 contacts supporting structural integrity

To shed light on why Toc75 and Toc33/34 are relatively more important for TOC structural integrity **(Fig. 2)**, and potentially also for the import mechanism, we further scrutinized the TOC-P structural model. Within the GTPase heterodimer of TOC-P, the two components are positioned around a two-fold rotational axis relative to each other **(Extended Data Fig. 3a)**. Two tight GTP binding pockets exist at the Toc33-Toc159 interface for which the prediction confidence is high **(Fig. 3b,c)**. In these pockets, Toc33-GTP2/GTP1 hydrogen-bond interactions (Toc33-GTP2: KSS [aa 49-51], H160, N209; Toc33-GTP1: R130) and Toc159-GTP1/GTP2 hydrogen-bond interactions (Toc159-GTP1: G865, GKSA [aa 867-870], S889, H982, N1036; Toc159-GTP2: D989) are at highly conserved positions **(Extended Data Fig. 3d,e)**. This implies that the two GTPs likely play a role in the dimerization process. Of these residues, Toc33-R130 was previously implicated in homodimerization in vitro, potentially acting as an arginine finger with a catalytic role in GTP hydrolysis^29–31^. In fact, the AF3-predicted GTPase heterodimer is similar overall to the crystal structure of a homodimer of *Pisum sativum* Toc34 formed in vitro, providing further strong support for the veracity of the AF3 model^30^ **(Supplementary Fig. 4)**.

As already noted, the GTPase heterodimer is connected to the top of the hybrid β-barrel with two linkers **(Fig. 3c)**. Linker-1 (Toc33, D251-G264) connects the GTPase domain and single transmembrane helix (TMH) of Toc33, the latter ending adjacent to the cytosolic face of the Toc75 β-barrel domain **(Extended Data Fig. 4a,b)**; notably, a highly-conserved proline kink (P269) at the cytosolic end of the Toc33 TMH may help in the orientation of the linker and GTPase domain towards the channel. Linker-2 (Toc159, R1078-P1097) brings the GTPase and membrane domains of Toc159 into close proximity **(Extended Data Fig. 4c,d)**. To analyse the flexibility of these linkers, we performed molecular dynamics (MD) simulations of the TOC complex embedded in a 1-palmitoyl-2-oleoyl-*sn*-glycero-3-phosphocholine (POPC) lipid bilayer **(Fig. 3h)**. The results revealed that linker-2 is more flexible than linker-1 **(Supplementary Videos 1 and 2, and Fig. 3i,j)**. As a consequence, the GTPase domain of Toc159 is more mobile than the GTPase domain of Toc33, showing larger root mean square deviation (RMSD) values **(Supplementary Videos 1 and 2, and Fig. 3i,j)**. Movement of the Toc159 GTPase domain is likely influenced by its α-2 helix through establishment of spontaneous membrane contacts **(Supplementary Fig. 5)**. Overall, this dynamic behaviour, which does not occlude the channel entry gate at the cytosolic side, may be relevant for preprotein hand-over from the receptor module to the channel module.

Another remarkable feature of the structure is that the membrane domain of Toc75 and the membrane domain (consisting of β-barrel and embedded segments) of Toc159 cooperate in the formation of a hybrid β-barrel comprising a total of 30 antiparallel β-strands: a 16-strand β-sheet from Toc75, and a 14-strand β-sheet from Toc159 **(Fig. 3c and Extended Data Fig. 5a-e)**. The two constituent β-sheet domains are joined by a β-seam, whereby the first β-strand (β1) of Toc75 is bonded to the last β-strand (β14) of Toc159 **(Fig. 3c and Extended Data Fig. 5c)**. Such interactions create a flat face on one side of the β-barrel. The C-terminal transmembrane helix of Toc159 traverses the outer face of the β-seam likely stabilizing the β-barrel. On the opposite side, the β-sheet domains of Toc75 (β13-16) and Toc159 (β1-4) converge by curling inward and closing off the β-barrel **(Extended Data Fig. 5d,e)**. This resembles the hybrid β-barrel of a BamA-substrate complex **(Extended Data Fig. 5f)**^32^. In addition, a part of an embedded segment of Toc159 extends across this junction and underneath the Toc75-side of the β-barrel **(Extended Data Fig. 5c,d)**. This overall arrangement of the hybrid β-barrel likely allows the protein import channel to expand and contract flexibly in order to accommodate (un)folded protein domains of different sizes^33,34^.

The Toc75 part of the dimeric β-barrel establishes contacts with the TMH of Toc33 and the TMH of Toc159 on different sides **(Fig. 3c and Extended Data Fig. 5e)**. Moreover, interface analysis of TOC-P identified Toc75 as a hub for interactions with Toc33 both in the membrane region and at the cytosolic side, and with Toc159 mainly at the membrane level **(Extended Data Fig. 6a,b)**. Indeed, a missense mutation (G518R of mature Toc75) in *kd-toc75* (*mar1*) affecting a conserved position that is proximal to these Toc75 contact sites destabilizes the TOC-P and TOC-N complexes **(Fig. 2 and Extended Data Fig. 6a,d)**, supporting their importance for complex integrity. In contrast, Toc33-Toc159 interactions are limited to the GTPase domains **(Fig. 3c and Extended Data Fig. 6c)**. While Toc33 is important for the stabilization of the holocomplex (most likely through its TMH and GTPase domains) **(Fig. 2c)**, it appears that the TMH of Toc159 is not. This is clear because the *kd-toc159* (*fts1*) mutant (which lacks the C-terminal TMH due to a premature stop at codon 1472) displays a drastic reduction in Toc159 accumulation without impacting levels of Toc75 and Toc33 accumulation **(Fig. 2c and Extended Data Fig. 7a)**. Moreover, Toc159 in *kd-toc159* but not that in wild type is destabilized by alkaline treatment of a membrane fraction, suggesting that the missing TMH is needed to properly anchor and stabilize the Toc159 β-barrel in the membrane **(Extended Data Fig. 7b)**. The results also imply that Toc75-Toc33 can assemble as a stable subcomplex without Toc159 **(Fig. 2c)**.

Wishing to assess the extent to which the protein import apparatus has been structurally conserved during chloroplast evolution, we compared the AF3-predicted TOC-P complex with the cryo-EM structures of TOC-TIC from the green alga *Chlamydomonas reinhardtii*^27,28^ **(Supplementary Fig. 6a)**. The latter lack information on the cytosolic GTPase domains of the receptors (termed Toc34 and Toc120/Toc90) **(Supplementary Fig. 6a,b)**, presumably due to the flexibility of the linkers described earlier. Thus, we applied AF3 to model the algal TOC complex, revealing that the missing domains assume a heterodimeric arrangement analogous to that in plants (ipTM = 0.75; pTM = 0.74) **(Supplementary Fig. 6c)**. Comparing the different structures, we found that TOC-P from *A. thaliana* resembles the cryo-EM and AF3-predicted structures of the algal TOC complex, indicating that structural organization of the TOC complex is broadly conserved in plants and green algae **(Supplementary Fig. 6a,c)**. However, affinity purified TOC-P complexes from our *HA-Toc75* line as well as from TAP-tagged transgenic lines (TAP-Toc159, TAP-Toc33) showed negligible interaction with inner membrane TIC components **(Extended Data Fig. 8)**. This revealed a clear and remarkable distinction from the algal TOC apparatus which is tightly integrated with the TIC apparatus in an interlocked TOC-TIC supercomplex^27,28^. This implies that the plant and algal protein import systems are substantially different.

### Toc75 is a key player in the biogenesis of TOC complexes

Our biochemical and structural analyses revealed differing contributions of the TOC components to overall integrity and architecture of the complex. To investigate if these differences are reflected also in the assembly pathway, we studied the biogenesis of the TOC complex. We began by analysing the expression of *TOC* genes during de-etiolation – the greening process in which chloroplast biogenesis results in photosynthetic establishment^35–37^. Dark-grown wild-type seedlings were exposed to long-day photoperiods for 7 days, during which time photosynthesis was gradually initiated. During the first 5 days of illumination, expression of the light-harvesting complex gene *LHCB1* increased rapidly and by an order of magnitude, as expected **(Fig. 4a)**. Interestingly, *TOC75* expression followed a very similar pattern of dramatic induction to *LHCB1*. In contrast, all other *TOC* genes were only moderately upregulated. These results suggested that *TOC75* expression plays a leading role during TOC biogenesis.

**Fig. 4.**
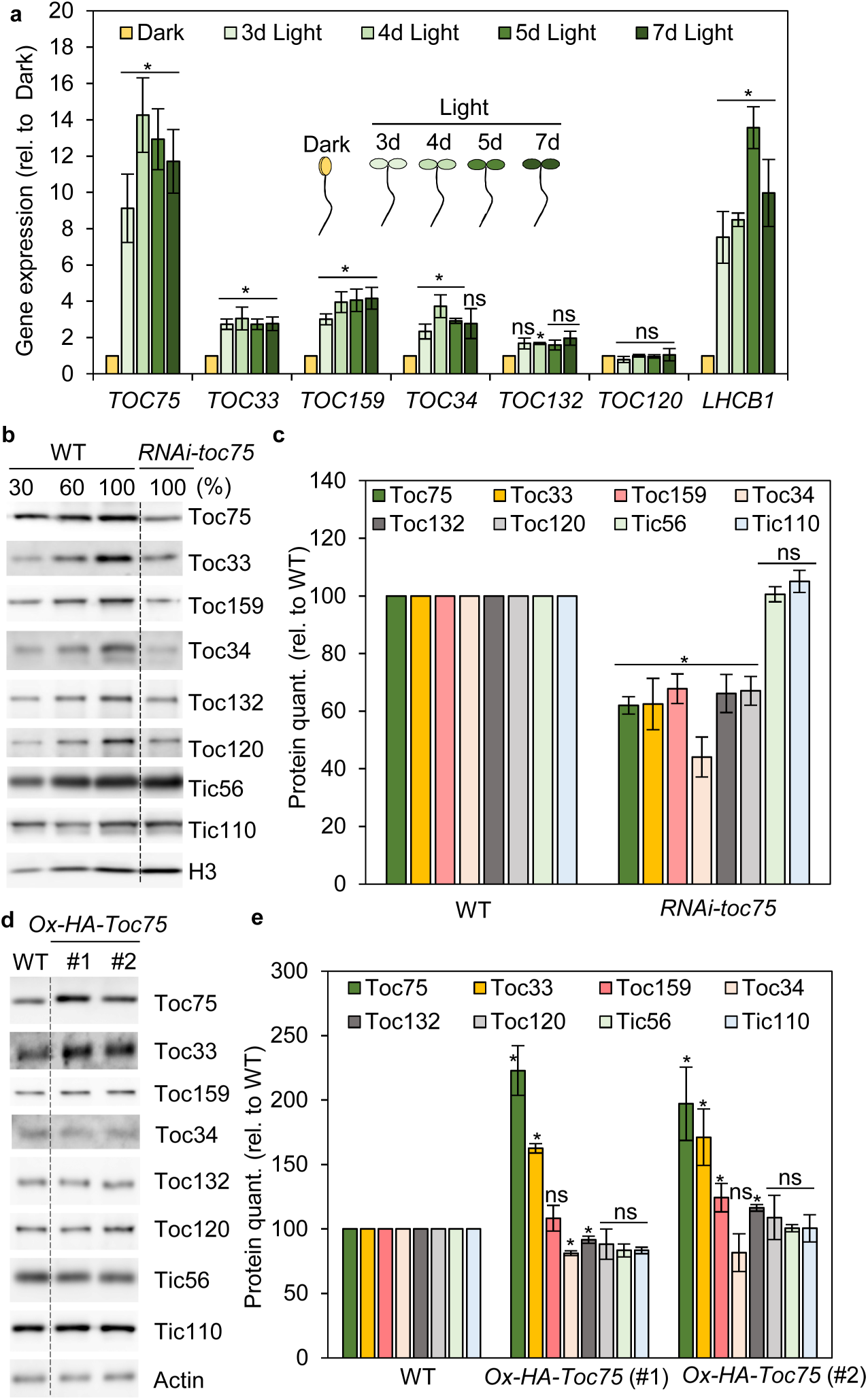
Identification of Toc75 as the central player in TOC complex formation. **a.** Gene expression analysis of TOC components during de-etiolation by qRT-PCR. Dark-grown WT seedlings were exposed to long-day cycles (16-h light / 8-h dark) for between 3 and 7 days (3d-7d Light). Expression data for genes of interest were normalized using data for *ACTIN2*. Asterisks indicate significance according to paired one-tailed Student’s t tests comparing the de-etiolation time points with the Dark sample (\**p* < 0.05; ns, not significant). **b,c.** Immunoblotting analysis of WT and *RNAi-Toc75* knockdown plants (**b**), and corresponding protein quantification by densitometry (**c**). Seedlings were grown for 10-12 days on medium containing 50 mM dexamethasone prior to analysis. **d,e.** Immunoblotting analysis of WT and *Ox-HA-Toc75* over-expressor plants (**d**), and corresponding protein quantification by densitometry (**e**). In **c** and **e**, asterisks indicate significance according to paired one-tailed Student’s t tests comparing *Ox-HA-Toc75* with WT (\**p* < 0.042; ns, not significant). All values are means ± SEM (n = 3 experiments).

To investigate whether *TOC75* gene expression changes can affect the accumulation of TOC components, we first studied an RNA interference (RNAi) line in which *TOC75* mRNA was reduced to ∼20% of the wild-type level^22^ **(Fig. 4b,c)**. Immunoblotting analysis showed that the reduced accumulation of Toc75 in the *RNAi-toc75* plants resulted in an overall reduction in the accumulation of all TOC-P and TOC-N components, while leaving TIC components unaffected. These results supported the view that *TOC75* expression can to some extent drive the accumulation of the TOC-P and TOC-N components. To further test this hypothesis, we employed two *TOC75* over-expression lines (*Ox-HA-Toc75*, 1 and 2) that were generated in the *kd-toc75* background which, as described earlier, selectively reduces TOC (not TIC) protein accumulation **(Fig. 2a)**. In these lines, the expression of *TOC75* was enhanced 5-8 fold relative to wild type, while the expression of *TOC33* and *TOC159* was unaltered **(Extended Data Fig. 9)**. Notably, immunoblotting revealed that all TOC-P and TOC-N proteins were recovered to at least wild-type levels in the *Ox-HA-Toc75* lines **(Fig. 4d,e)**. In fact, Toc33 protein even exceeded the wild-type level despite *TOC33* transcript abundance being unchanged. Thus, on the basis of all these results, we concluded that Toc75 is a leading and central player in the biogenesis of TOC complexes.

### Toc75 synthesis precedes that of Toc159 in the TOC assembly process

To shed further light on the synthesis and assembly of TOC complexes, wild-type and *HA-Toc75* seedlings were subjected to in vivo radiolabelling using inorganic sulphate (Na_2_^35^SO_4_) over a time-course^38^. Very young seedlings were employed so as to capture an active phase of photosynthetic establishment. Total radiolabelled seedling extracts were solubilized and subjected to anti-HA affinity purification, and the purified complexes were analysed by SDS-PAGE and autoradiography **(Fig. 5a)**.

**Fig. 5.**
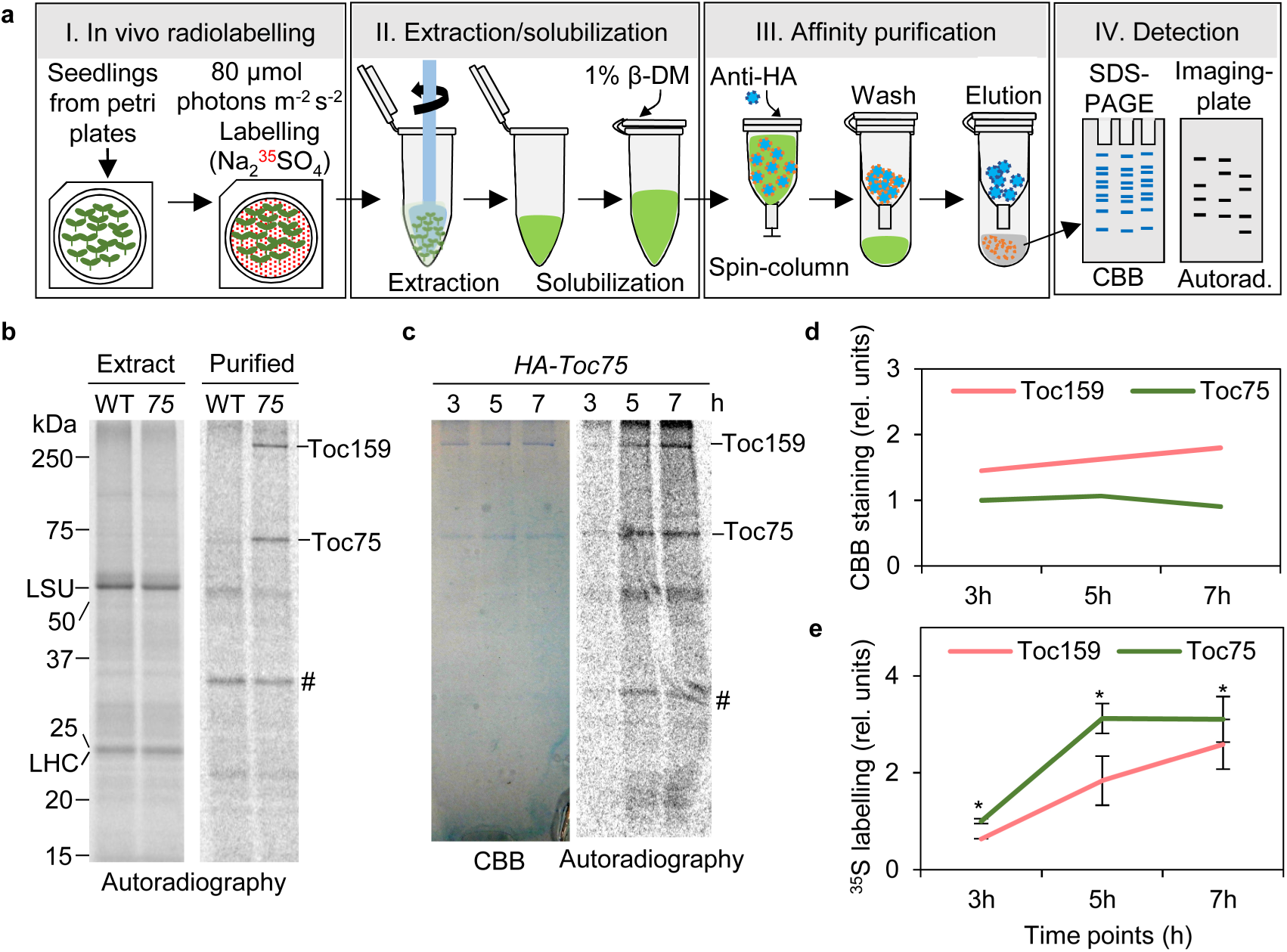
In vivo labelling analysis reveals the order of synthesis of TOC components. **a.** Schematic summary of the experimental procedure. **b.** Radiolabelling of 7-8-day-old WT and *HA-Toc75* (*75*) transgenic plants with inorganic sulphate (Na_2_^35^SO_4_) for 16 h. Labelled protein samples extracted from the plants were subjected to HA-affinity purification and/or analysis by SDS-PAGE and autoradiography. Samples were solubilized with 1% β-DM before affinity purification. Total protein extracts (Extract) and affinity-purified samples (Purified) are shown. Positions of molecular weight markers (sizes in kDa), and two prominent chloroplast proteins (LSU, LHC), are indicated at left. A non-specific band is indicated (#). **c.** Similar radiolabelling of HA-Toc75 plants for shorter time periods. Affinity purified samples from 3, 5 and 7 h labelling time points were prepared as in **b**. Resolved polypeptides in the purified samples were visualized by either staining (Coomassie brilliant blue, CBB) or autoradiography. A non-specific band is indicated (#). **d,e.** The CBB (**d**) and autoradiography (**e**) signals from **c** were quantified by densitometry. Asterisks indicate significance according to paired one-tailed Student’s t tests comparing the labelling of Toc159 with the labelling of Toc75 (\**p* < 0.015). All values in **e** are means ± SEM (n = 2-3 experiments). In **e**, the labelling of Toc159 is divided by a factor of 1.25 (because the cysteine + methionine residue counts for Toc75 and Toc159 are 20 and 26, respectively).

After an extended labelling reaction (16 h), the labelling intensities of two chloroplast diagnostic proteins, Rubisco large subunit (LSU) and light-harvesting complex protein (LHC), were identical in the two genotypes, indicating efficient labelling of the chloroplasts **(Fig. 5b)**. However, the TOC complex could only be purified from the *HA-Toc75* line, as expected. The purified complex showed strong labelling of Toc75 and Toc159 (Toc33, which has a low cysteine/methionine content (9 residues), was unfortunately obscured by a comigrating nonspecific band), indicating that these components were undergoing active synthesis. Although the other TOC components (i.e., TOC-N components) were shown to be present in comparable samples by immunoblotting **(Fig. 1f)**, their labelling was not detected.

Next, to reveal the precise order of synthesis, we analysed TOC complexes from shorter labelling reactions (3 to 7 h) **(Fig. 5c)**. Based on the staining intensity (by Coomassie brilliant blue, CBB) of Toc159 and Toc75, the purified sample contained similar yields of Toc75 and Toc159 (staining was slightly higher for Toc159, likely because of its larger size) **(Fig. 5c,d)**. However, the Toc75 protein was more rapidly labelled. Labelling of Toc75 was readily detectable from 3 h, and essentially plateaued by 5 h of labelling **(Fig. 5c,e)**. The labelling pattern of Toc159 was similar, but progressed more slowly and was still rising at 7 h, never quite reaching the levels seen in Toc75 (after adjusting for the cysteine/methionine contents of the proteins: 26 in Toc159, 20 in Toc75) **(Fig. 5e)**. These findings reveal that Toc75 is synthesized just before Toc159, likely as part of a step-wise assembly process. Unfortunately, it was not possible to follow Toc33 labelling in this experiment (as noted earlier). However, our biochemical data **(Fig. 2)** and structural modelling **(Fig. 3)** showed that Toc33 is required for the structural integrity of the TOC-P complex. We therefore infer that Toc33 is integrated just after the integration of Toc75, yielding a stable intermediate complex, Toc75-Toc33, that rapidly assembles with Toc159 to form the mature TOC-P translocon in chloroplasts **(Figs. 5d and 6)**.

**Fig. 6.**
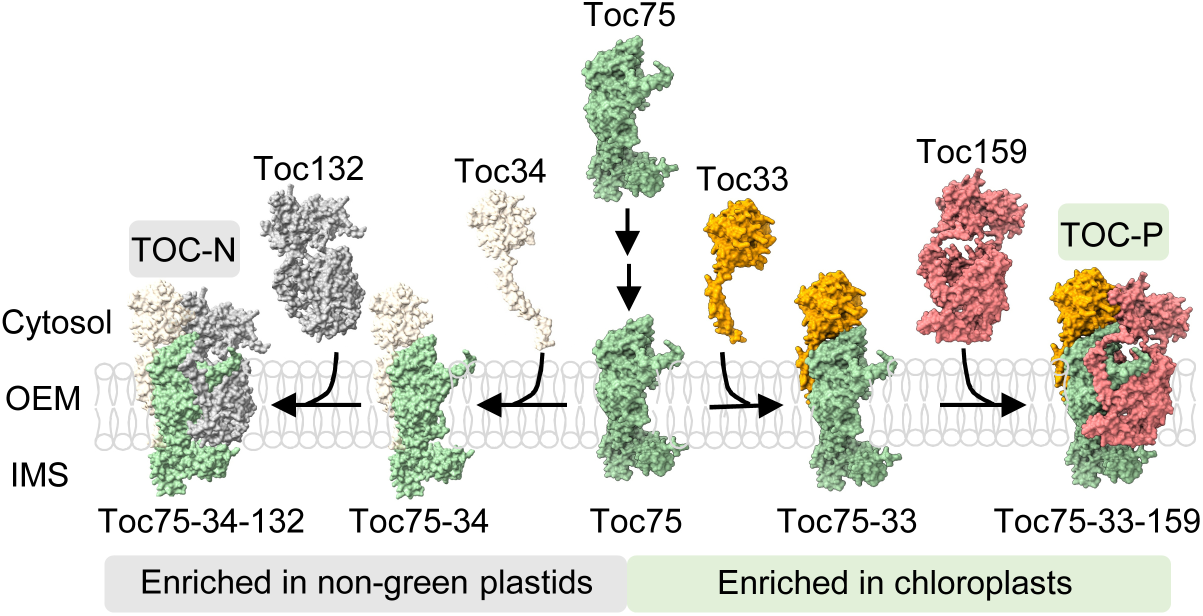
Model for the step-wise assembly of TOC complexes. Assembly is initiated by gene expression of *TOC75*; and upon translation, Toc75 is integrated into the outer envelope membrane of the organelle. Newly inserted Toc75 is stabilized by the integration of Toc33, forming a mutually-stabilized Toc75-Toc33 assembly intermediated complex. In the next step, in chloroplasts, Toc75 rapidly assembles with Toc159 to yield the mature TOC-P translocon complex. Based on our genetic, biochemical and structural modelling data, we infer that Toc75 assembles sequentially with Toc34 and Toc132 (or the redundant Toc120) in similar fashion to form TOC-N complexes; this is a minor process in chloroplasts where TOC-P predominates, but is the dominant process in non-green plastids such as leucoplasts where TOC-N is the major complex.

Although the TOC-N components were undetectable by either staining or radiolabelling in these experiments (reflecting the fact that TOC-P is the dominant configuration in photosynthetic tissues), based on our preceding analyses **(e.g., Figs. 1, 2, 4 and Extended Data Fig. 6)**, we further infer that TOC-N biogenesis follows a similar pathway **(Fig. 6)**.

### Receptor mutant analyses reveal functional differences between TOC-P and TOC-N

Having elucidated compositional and structural differences between the TOC-P and TOC-N complexes, we next wished to assess for functional differences. To this end, we employed a microscopy-based ratiometric approach to assess protein import in the *ko-toc33* and *ko-toc34* mutants which lack characteristic receptors of TOC-P and TOC-N, respectively. To create model clients for these assays, the transit peptides (TP) of Rubisco small subunit precursor (SSU; a photosynthetic preprotein) and pyruvate dehydrogenase E1α subunit precursor (E1α; a non-photosynthetic preprotein)^15,39^ were fused with either YFP or CFP, yielding four fusions as follows: TP-SSU-YFP, TP-E1α-YFP, TP-SSU-CFP and TP-E1α-CFP.

First, these fusions were assessed in wild-type protoplasts to verify that their behaviour was as expected. After their expression in different pairwise combinations, the transfected cells were analysed by confocal microscopy. Although CFP fluorescence is inherently weaker than that of YFP, the TP-SSU-CFP fusion generally produced stronger chloroplast-associated fluorescence than TP-E1α-YFP in co-transfected cells, resulting in low YFP/CFP ratios **(Extended Data Fig. 10a,b)**. In contrast, in reciprocal experiments using TP-SSU-YFP and TP-E1α-CFP, an opposite YFP/CFP ratio trend was seen **(Extended Data Fig. 10b)**. Importantly, the observed fluorescence intensity differences were not due to differences in expression, as revealed by RT-PCR analysis **(Extended Data Fig. 10c)**. These data indicated that the SSU TP was more effective at delivering protein import into photosynthetic chloroplasts, which is as expected. Therefore, this ratiometric system was implemented to analyse protoplasts from *ko-toc33* and *ko-toc34* mutants, to test for specific effects of the mutations on the efficacy of the SSU and E1α TPs. In assays with TP-E1α-YFP and TP-SSU-CFP, higher YFP/CFP ratio values were seen in *ko-toc33* than in *ko-toc34* **(Fig. 7a,b [left side of b])**; whereas in the reciprocal experiment using SSU-YFP and E1α-CFP, the opposite trend was observed **(Fig. 7b, right side)**.

**Fig. 7.**
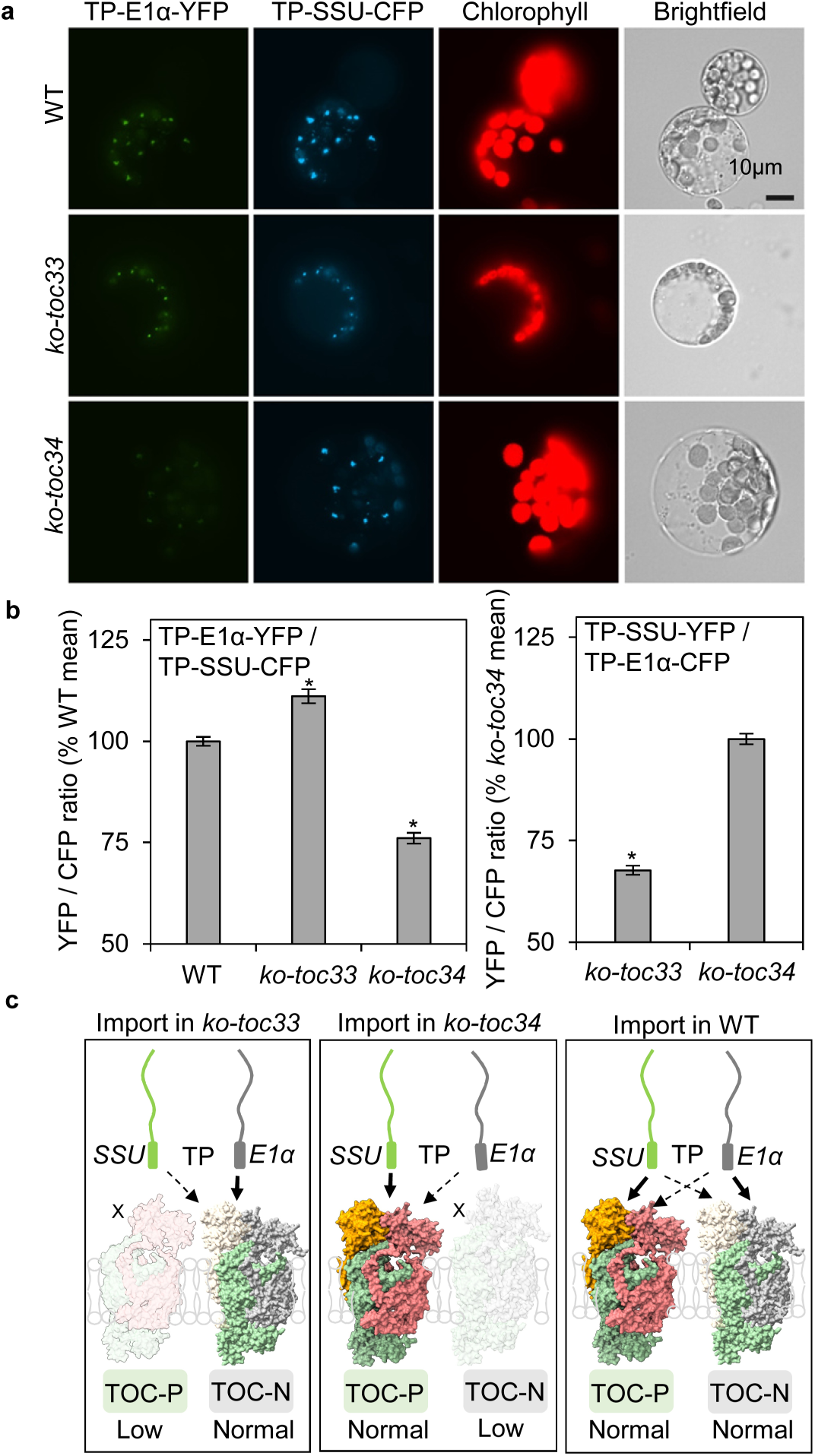
In vivo protein import analyses reveal client specificity differences between *ko-toc33* and *ko-toc34* mutants. **a,b.** Ratiometric analysis of pairs of fusion proteins, each comprising a transit peptide (TP) of Rubisco small subunit (SSU) and TP of pyruvate dehydrogenase E1α subunit (E1α) linked to YFP or CFP. Protoplasts from WT, *ko-toc33* and *ko-toc34* plants were co-transfected with plasmids encoding the indicated TP fusions, and fluorescence microscopy was used to detect the transiently-expressed proteins. Qualitative analysis of the relative chloroplast-targeting efficiencies of the TP-E1α-YFP and TP-SSU-CFP fusions is shown (**b**). The YFP (green; left panels) and CFP (cyan; centre-left panels) fluorescence signals were recorded, alongside chlorophyll autofluorescence (red; centre-right panels) and brightfield images (right panels). Each set of images is of the same cell, and is representative of a large number of images derived from four independent experiments. The YFP/CFP fluorescence intensity ratios within individual chloroplasts were quantified (**b**), both for the experiment shown in **a** (left graph) and for the inverse experiment (right graph). For each genotype, the data shown comprise ∼1000-2000 individual ratio measurements, derived from a total of up to 66 different protoplasts from four separate experiments. Asterisks indicate significance according to two-tailed Student’s t tests comparing the mutants with WT (\**p* < 0.001). All values are means ± SEM (n = 904-1865). **c.** Schematic representation of the protein import properties of the *ko-toc33* and *ko-toc34* mutants, and of WT. Import of TP-SSU preprotein (a photosynthetic client) is reduced in *ko-toc33*, whereas import of TP-E1α preprotein (a non-photosynthetic client) is reduced in *ko-toc34*. Thus, the SSU and E1α preproteins are preferentially imported by TOC-P and TOC-N, respectively, in the WT. Calculated pI values of the TPs of SSU and E1α are substantially different (10.05 and 12.13, respectively); this suggests that charge differences between TPs, together with surface electrostatic potential differences between TOC-P and TOC-N **(see Extended Data Fig. 2f)**, may underly the observed client preferences.

Overall, these data provide in vivo support for the hypothesis that TOC-P and TOC-N are functionally distinct, in accordance with their respective predominance in photosynthetic and non-photosynthetic tissues, owing to their possession of receptors with specificity for different transit peptides **(Fig. 7c)**.

## DISCUSSION

The protein translocon in the outer membrane of chloroplasts and other plastids, TOC, comprises an Omp85-type β-barrel channel, Toc75, and two related GTPase receptors, Toc33 and Toc159^1,3,4,6^. Because the receptors are encoded by small gene families in plants, it has long been proposed that different TOC systems may exist under different circumstances^3^. However, while evidence for functional differences among the various receptor isoforms has been presented, whether or not these isoforms indeed assemble selectively into distinct TOC complexes, for example in different plastid types^11,13,14^ has remained unclear^16,40^. A major challenge in this area has been the low abundance of the TOC machinery. We circumvented such problems by employing transgenic plants expressing tagged TOC components to enable efficient affinity-purification strategies even from non-photosynthetic tissues. This formed just one strand of a heavily integrated approach (combining diverse experimental approaches with advanced structural analysis) that enabled us to detail the assembly of two structurally- and functionally-distinct TOC complexes, termed TOC-P and TOC-N, that predominate in photosynthetic and non-photosynthetic tissues, respectively.

After applying biochemistry and genetics to elucidate subunit interactions within the TOC-P and TOC-N complexes, we implemented an integrative structural approach (comprising AF3, negative staining EM, XL-MS and MD) to uncover architectural features of the complexes, and the differences between them. This analysis revealed a novel heterodimeric-GTPase– heterodimeric-β-barrel arrangement, with the two heterodimeric modules being coupled by a pair of linkers **(Fig. 3)**. In TOC-P, the GTPase module comprises Toc33 and Toc159 whereas the β-barrel module comprises Toc75 and Toc159. The latter module is well-aligned with recently-published cryo-EM structures of TOC-TIC supercomplexes from the green alga *C. reinhardtii*^27,28^. However, the published cryo-EM structures completely lacked information on the receptor GTPase domains, presumably due to the way they are flexibly linked to the β-barrel module, as our results reveal. This highlights a key advantage of the integrative approach employed here over solely cryo-EM-based approaches. We speculate that flexibility is a key feature of the TOC machine, enabling the receptor GTPase module to flip-flop back and forth so as to collect precursor protein clients in the cytosol and then deliver them into the β-barrel channel module. The TOC GTPase heterodimer calls to mind another interaction between related GTPases, between SRP54 and its membrane receptor FtsY, which occurs during nascent protein delivery to the bacterial cell membrane^41^. However, in that case the association is only transient in nature, whereas the TOC GTPases appear stably associated.

Even though the TOC β-barrel module is similarly arranged in plants and algae, our results show that the two systems have nonetheless diverged functionally. Whereas in algae the TOC is deeply and stably integrated with the TIC machinery^27,28^, we find that in plants the two translocons are readily separable, a result that aligns well with recent cryo-EM structures of the plant TIC and an associated Ycf2-FtsHi motor complex which lack TOC components^42^. This feature may be a special adaptation in plants to enable greater flexibility in the import system. One can easily envision how it may be crucial to have much more dynamic TOC-TIC associations in plants, to enable full exploitation of the advantages that having multiple TOC subtypes can bring, for example when protein import requirements change dramatically during development. This dynamism might be especially important during developmental phases in which the plastids change type via organellar proteome remodelling (such as in de-etiolation or fruit ripening), and it may be facilitated by the proteolytic removal of unneeded TOC components^1,20,43^. Implicit in this hypothesis is a need for variant TOCs that differ functionally, and our data clearly indicate that this requirement is met. The TOC-P and TOC-N complexes have structural differences, including surface electrostatic potential differences at the entry side of the complex, as well as functional differences in relation to preprotein client specificity **(Fig. 7 and Extended Data Fig. 2)**.

On consideration of our in vivo labelling results in conjunction with the genetics, expression and structural data, we proposed a model for TOC biogenesis **(Fig. 6)**. In this, new TOC complexes are formed in a co-assembly process in which Toc75 plays a driving role. Synthesis of Toc75 primes a new complex, which is then stabilized in the membrane by the addition of Toc33 before final completion with the insertion of Toc159 **(Fig. 6)**. This model is consistent with previous in vitro reconstitution experiments conducted using isolated chloroplasts or proteoliposomes^44–46^. Those studies showed that resident Toc33 and Toc75 are required to integrate Toc159 into the membrane, although the order in which Toc33 and Toc75 themselves are assembled was not addressed. It was further suggested that a heterodimeric interaction of the Toc159 GTPase domain with that of Toc33 promotes the integration of the Toc159 membrane-domain, revealing an additional parallel with our results. Other studies showed that resident Toc75 is sufficient to mediate the insertion of outer membrane proteins with similar characteristics to the Toc33 component^47^, which is again consistent with our TOC biogenesis model **(Fig. 6)**.

It is also informative to consider TOC biogenesis from an evolutionary perspective. As a member of the Omp85 superfamily of proteins^48^, it is clear that Toc75 is of prokaryotic origin and most likely derived from a cyanobacterial Omp85-type protein during endosymbiosis. Thus, we may infer that it was the first component to be recruited to the nascent protein import system of the chloroplast outer envelope membrane during the evolution of photosynthetic eukaryotes^49^. The presence of eukaryotic-type GTPase domains in the receptors, Toc33 and Toc159, led to the further conclusion that these components are of eukaryotic origin and were therefore added later^50^. The large receptor, Toc159, further evolved with the addition of terminal extensions at both ends (the N-terminal acidic domain, and the C-terminal membrane domain), delivering new capabilities related to client selectivity and channel flexibility. Interestingly, the Toc159 membrane β-barrel domain shares considerable similarities with the mitochondrial protein import channel, Tom40 (as well as with the related metabolite channel, VDAC)^51–53^, pointing to a remarkable evolutionary link between the protein import systems of the two prevalent endosymbiotically-derived organelles in eukaryotic cells **(Supplementary Fig. 7)**. In any case, it is clearly apparent that our TOC assembly model (in which Toc75 plays a leading role) is well aligned with these evolutionary aspects.

In conclusion, we have defined the properties of two distinct TOC complexes in plant plastids. The TOC-P and TOC-N complexes are structurally and functionally distinct, and are differentially synthesized and assembled to meet differing protein import demands in different plastid types. Furthermore, we show that TOC biogenesis is a dynamic and hierarchical process, in which the channel component Toc75 plays a central and driving role. Future work should seek to identify auxiliary factors that support the integration of new components during TOC complex assembly, in order to further elucidate the biogenesis mechanism. Moreover, our results suggest that fine-tuning the expression of Toc75 has the potential to enhance TOC accumulation, protein import, and possibly even photosynthesis, suggesting novel strategies for crop improvement.

## EXTENDED DATA FIGURES

**Extended Data Fig. 1.**
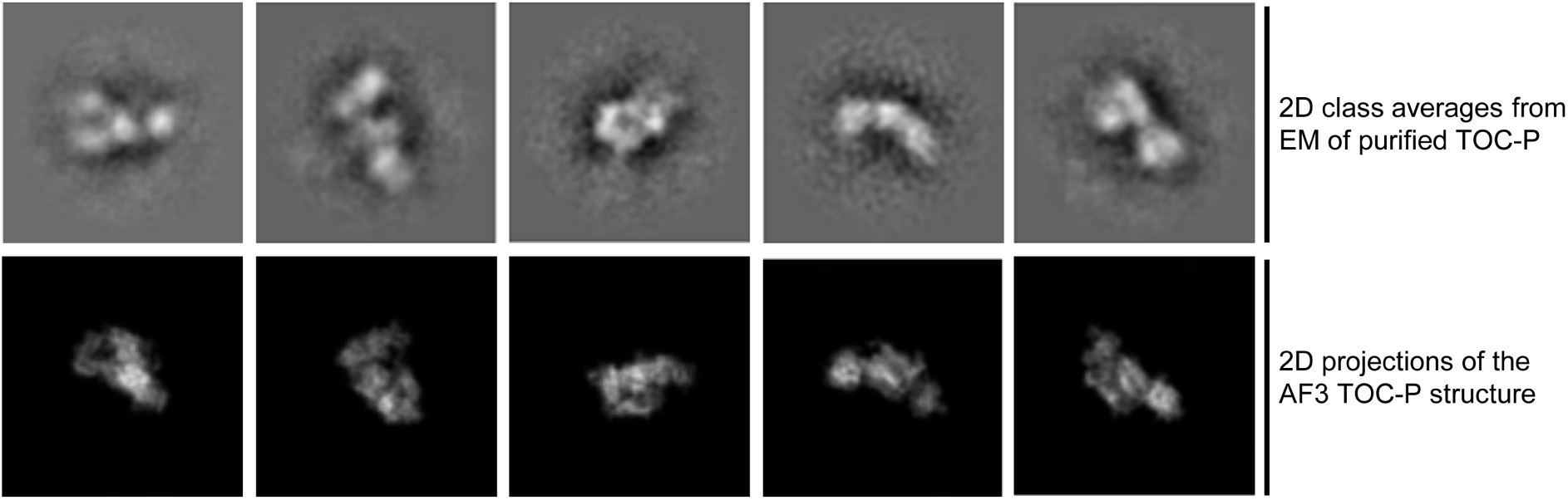
Negative staining EM analysis of the purified TOC-P complex. The affinity purified TOC-P complex was separated by size exclusion chromatography and the resulting peak fraction was analysed by negative staining and EM. Single particles from the EM images were picked and used to generate 2D class averages. In parallel, 2D projections of the AF3 TOC-P model **(see** Fig. 3) were generated by CryoSPARC, and these were compared with the 2D class average images from the EM analysis.

**Extended Data Fig. 2.**
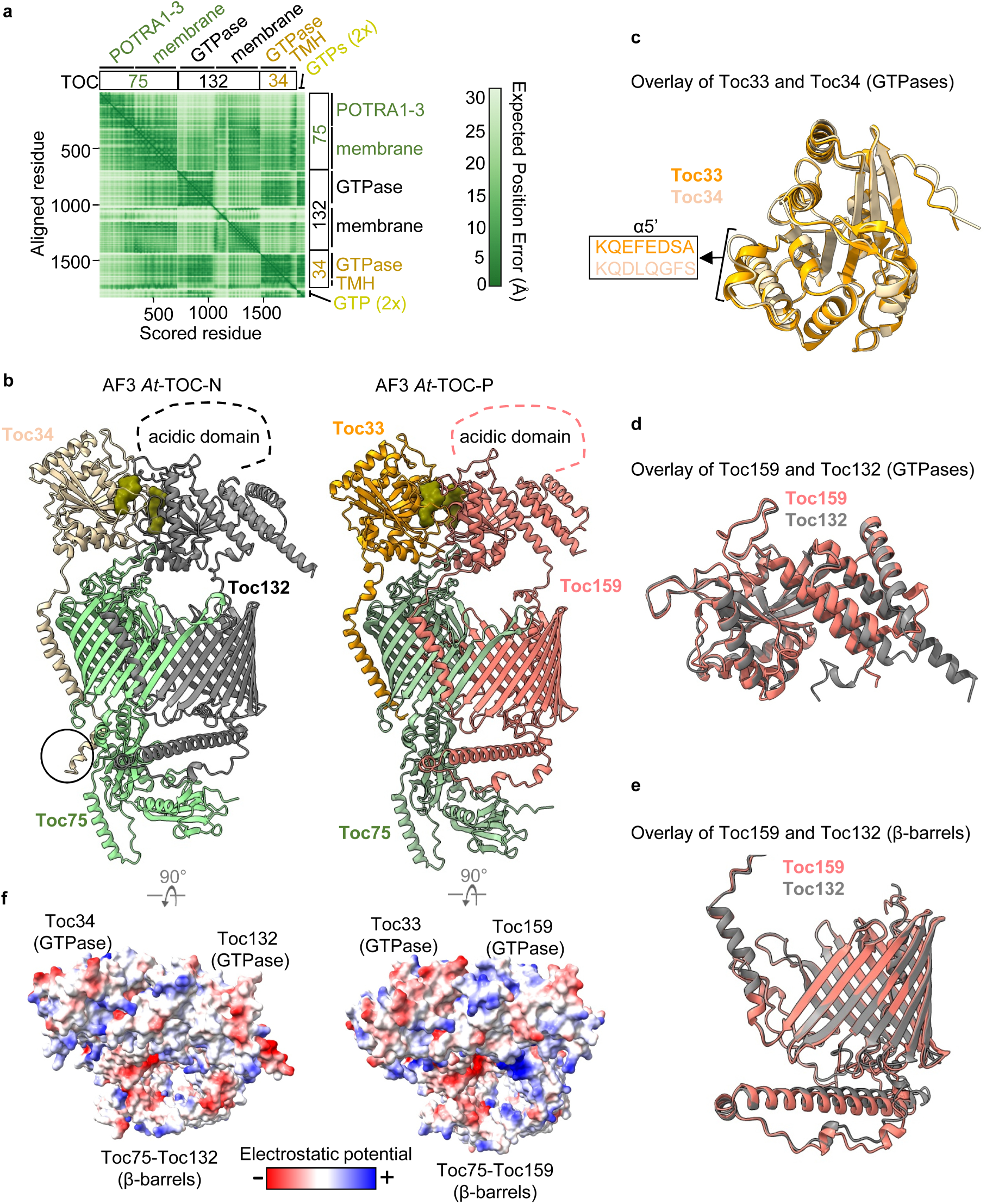
AF3 analysis of TOC-N reveals similarities to, and differences from, TOC-P. **a.** Predicted Alignment Error (PAE) plot from the AF3 analysis of TOC-N components and two GTPs. This analysis was conducted in exactly the same manner as the similar analysis of TOC-P **(see** Fig. 3). The sequences of Toc75 and Toc132 were N-terminally truncated to remove transit peptide and disordered A-domain sequences, respectively; the submitted sequences were 678 (Toc75), 720 (Toc132), and 313 (Toc34) residues long. **b.** Predicted structure of the TOC-N complex from AF3 (left side), alongside the similarly-generated TOC-P complex structure **(see** Fig. 3c) for comparison (right side). The circle shows an extended helix at the C-terminus of Toc34. Positions of the acidic A-domains of Toc159 and Toc132 (which were excluded because of their intrinsically disordered nature) are shown by the dashed lines. The two GTPs (depicted as surface models) are shown in olive green. **c.** Superimposition of the predicted structures of the GTPase domains of Toc33 and Toc34 (rotated 90° relative to panel **b**). The assignment of α5’ was according to a published component structure (PDB: 1H65)^30^. **d.** Superimposition of the predicted structures of the GTPase domains of Toc159 and Toc132. **e.** Superimposition of the predicted structures of the β-barrel domains of Toc159 and Toc132. **f.** Surface electrostatic potential representations of the TOC-N and TOC-P structures. Both images are top-down views from the cytosol. TOC-P (right) exhibits more acidic (negatively charged) patches on Toc33 and Toc75, and more basic (positively charged) patches on Toc159; in other words, Toc33 is more acidic than Toc34, and Toc159 is more basic than Toc132. These differences may play a role in determining the different client specificities of TOC-P and TOC-N by correlating with charge differences between preproteins. Calculated pI values of the transit peptides of the two preproteins assessed in this study (SSU and E1α) **(see** Fig. 7) are 10.05 and 12.13, respectively.

**Extended Data Fig. 3.**
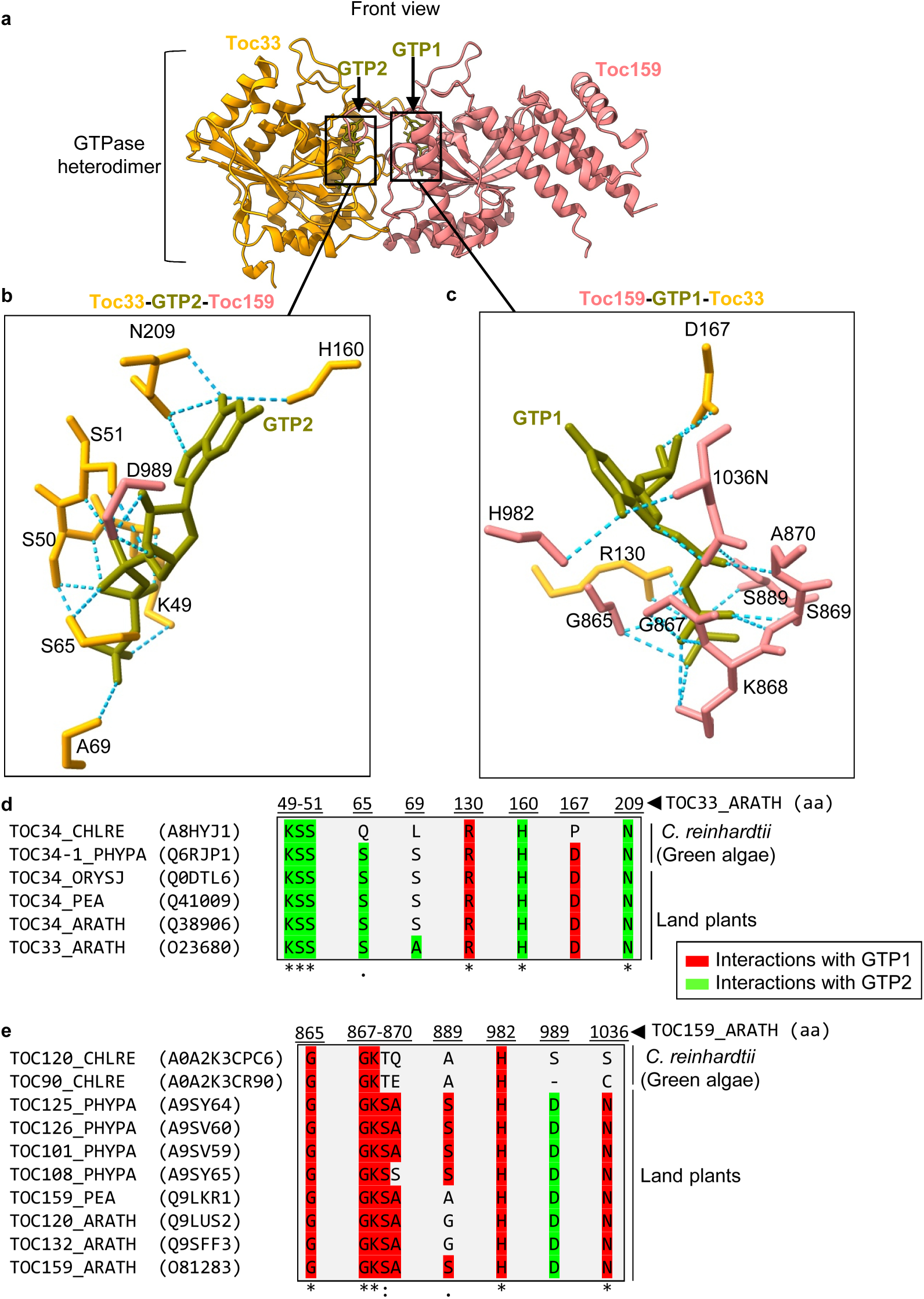
Two GTP-binding pockets exist at the interface of the GTPase heterodimer formed between Toc33 and Toc159. **a.** Structure of the GTPase heterodimer (front view), showing how the two GTPs lie at the interface of the dimer. GTP2 is principally associated with Toc33, whereas GTP1 is principally associated with Toc159. Boxes show the two GTP-binding pockets. **b,c.** Hydrogen-bond interactions in the two GTP-binding pockets. Interaction between Toc33, GTP2 and Toc159 (**b**), and between Toc159, GTP1 and Toc33 (**c**) are shown by the blue dotted lines. Both GTPases contribute to the binding of each GTP. **d,e.** Multiple sequence alignments (MSAs) showing conservation of the GTP-interacting amino acids of the receptor GTPases. Toc33-sequences were obtained from Uniprot for *C. reinhardtii* (A8HYJ1, TOC34_CHLRE), *Physcomitrium patens* (Q6RJP, TOC34-1-PHYP), *A. thaliana* (O23680, TOC33_ARATH; Q38906, TOC34_ARATH), *Oryza sativa* (Q0DTL6, TOC34_ORYSJ), and *P. sativum* (Q41009, TOC34_PEA) (**d**). Toc159-related sequences were obtained from Uniprot for *A. thaliana* (Q6S5G3, TOC90_ARATH; O81283, TOC159_ARATH; Q9LUS2, TOC120_ARATH, Q9SFF3, TOC132_ARATH), and *P. patens* (A9SY64, TOC125_PHYPA; A9SV60, TOC126_PHYPA; A9SV59, TOC101_PHYPA; A9SY65, TOC108_PHYPA) (**e**). Complete sequences were aligned in each case, but for simplicity only GTP-interacting amino acids are shown here (red, GTP1; green, GTP2). All residues shown in **d** and **e** are proximal to the interface of the GTPase heterodimer. Clustal Omega was used to perform both MSAs^54^, and the GTP-interacting residues were copied out manually. Coordinates at the top show positions in the *Arabidopsis* Toc33 (**d**) and Toc159 (**e**) proteins, corresponding to residues marked in **b** and **c**, respectively. The symbols at the bottom (*:.) indicate the degree of conservation at each position.

**Extended Data Fig. 4.**
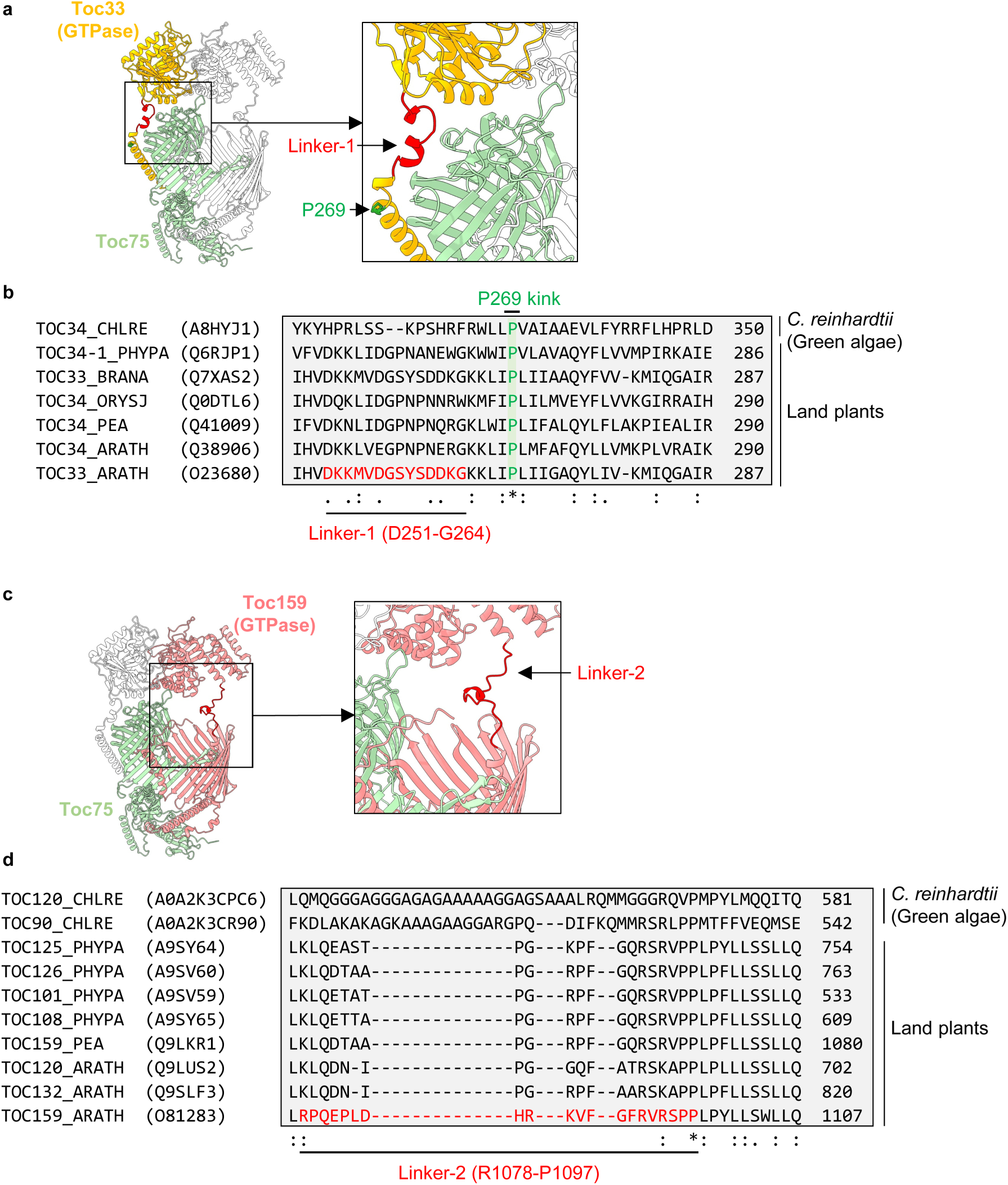
Coupling of the GTPase heterodimer with the β-barrel heterodimer is via two linkers. **a,c.** Elevated front views of the TOC-P structure showing the positions of linker-1 (**a**) and linker-2 (**c**). Linker-1 (D251-G264 of Toc33) positions the Toc33 GTPase domain at the cytosolic face of the β-barrel domain of Toc75. Linker-2 (R1078-P1097 of Toc159) positions the Toc159 GTPase domain close to the β-barrel domain of Toc159. The insets in each case show the linkers at higher magnification; a proline kink (P269) at the end of the Toc33 TMH is highlighted (**a, inset**). **b,d.** Multiple sequence alignments (MSAs) showing conservation of the two linkers. Toc33-related sequences were obtained from Uniprot for *C. reinhardtii* (A8HYJ1, TOC34_CHLRE), *P. patens* (Q6RJP, TOC34-1_PHYPA), *A. thaliana* (O23680, TOC33_ATHAL; Q38906, TOC34_ATHAL), *Brassica napus* (Q7XAS2, TOC33_BRANA), *O. sativa* (Q0DTL6, TOC34_ORYSJ), and *P. sativum* (Q41009, TOC34_PEA) (**b**). Toc159-related sequences were obtained from Uniprot for *C. reinhardtii* (A0A2K3CPC6, TOC120_CHLRE; A0A2K3CR90, TOC90_CHLRE), *A. thaliana* (O81283, TOC159_ARATH; Q9LUS2, TOC120_ARATH; Q9SLF3, TOC132_ARATH), *P. patens* (A9SY64, TOC125_PHYPA; A9SV60, TOC126_PHYPA; A9SV59, TOC101_PHYPA; A9SY65, TOC108_PHYPA) (**d**). Complete sequences were aligned in each case, but for simplicity only linker-proximal regions are shown here. Clustal Omega was used to perform both MSAs^54^, and the relevant regions were copied out manually. The sequences of the linkers in *Arabidopsis* Toc33 and Toc159 are highlighted in red, as is the position corresponding to the conserved Toc33 P269 residue (kink). Coordinates at right show positions in each protein sequence; symbols at the bottom (*:.) indicate the degree of conservation at each position. From **d**, it is evident that in land plants linker-2 is reasonably well conserved, but in green algae it contains sizeable alanine-rich insertions.

**Extended Data Fig. 5.**
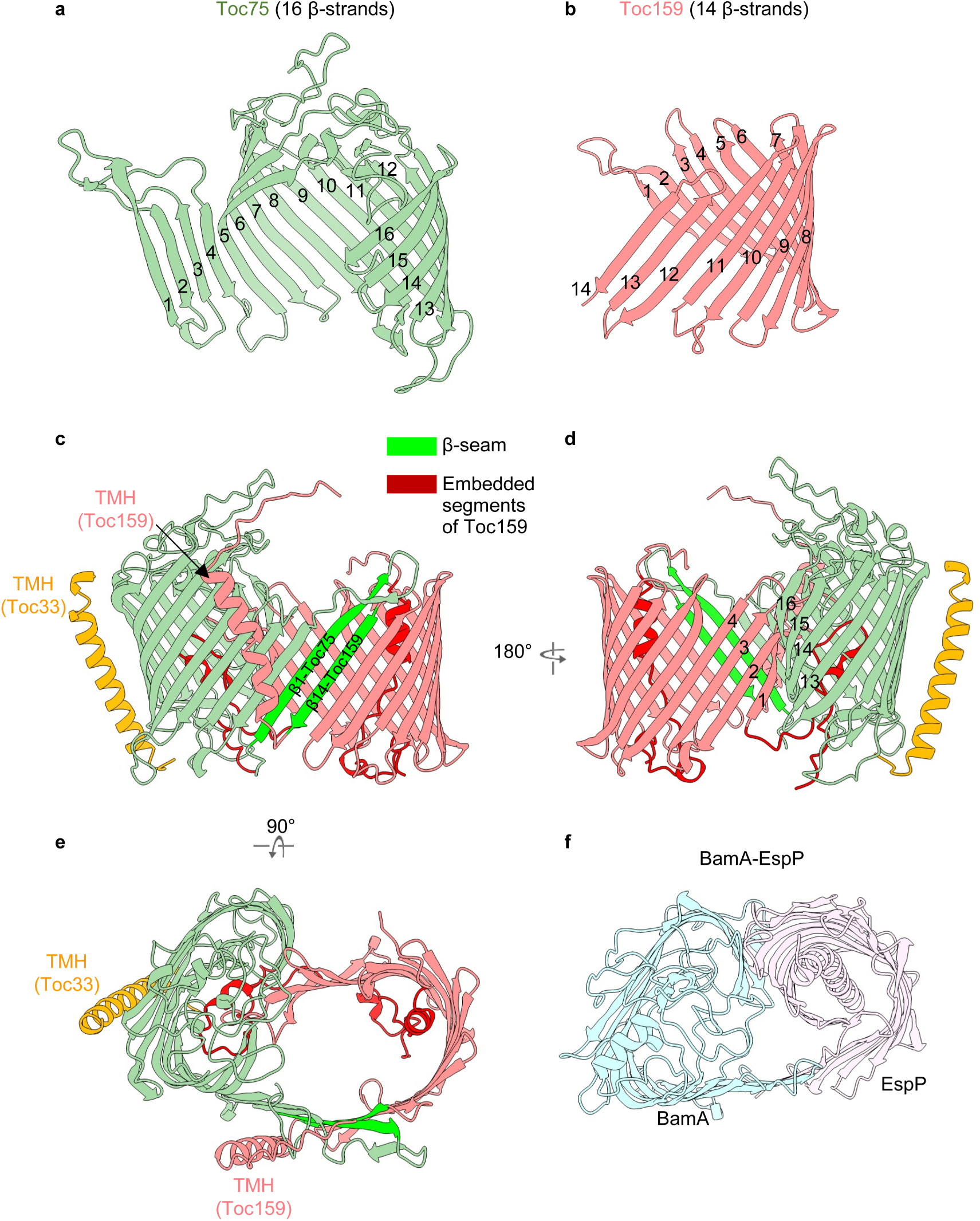
Structural organization of the heterodimeric β-barrel. **a,b.** Structures of the Toc75 and Toc159 β-barrel domains (each taken from the TOC-P complex model) are shown in isolation, to more clearly reveal the arrangement of the individual β-strands (16 in Toc75, 14 in Toc159). The images shown are front view (Toc159) or 90° rotation of front view (Toc75). **c.** At the front of the hybrid barrel, the β-sheet domains of Toc75 and Toc159 are connected by a β-seam between β1 of Toc75 and β14 of Toc159. **d.** At the other side of the barrel (back), the β-sheet domains of Toc75 (strands β13-β16) and Toc159 (strands β1-β4) curl inwardly to close the barrel. **e.** Top-down view of the hybrid barrel showing how the transmembrane helices (TMH) of Toc33 and Toc159 are in close proximity to the barrel on two faces of the complex. The Toc159 TMH traverses the β-seam and potentially acts to stabilize the hybrid barrel. **f.** For comparison, the published structure of a hybrid β-barrel assembly complex formed between bacterial BamA and its substrate protein EspP (PDB: 8BNZ)^32^. This is structurally similar to the Toc75-Toc159 hybrid barrel shown in **e**.

**Extended Data Fig. 6.**
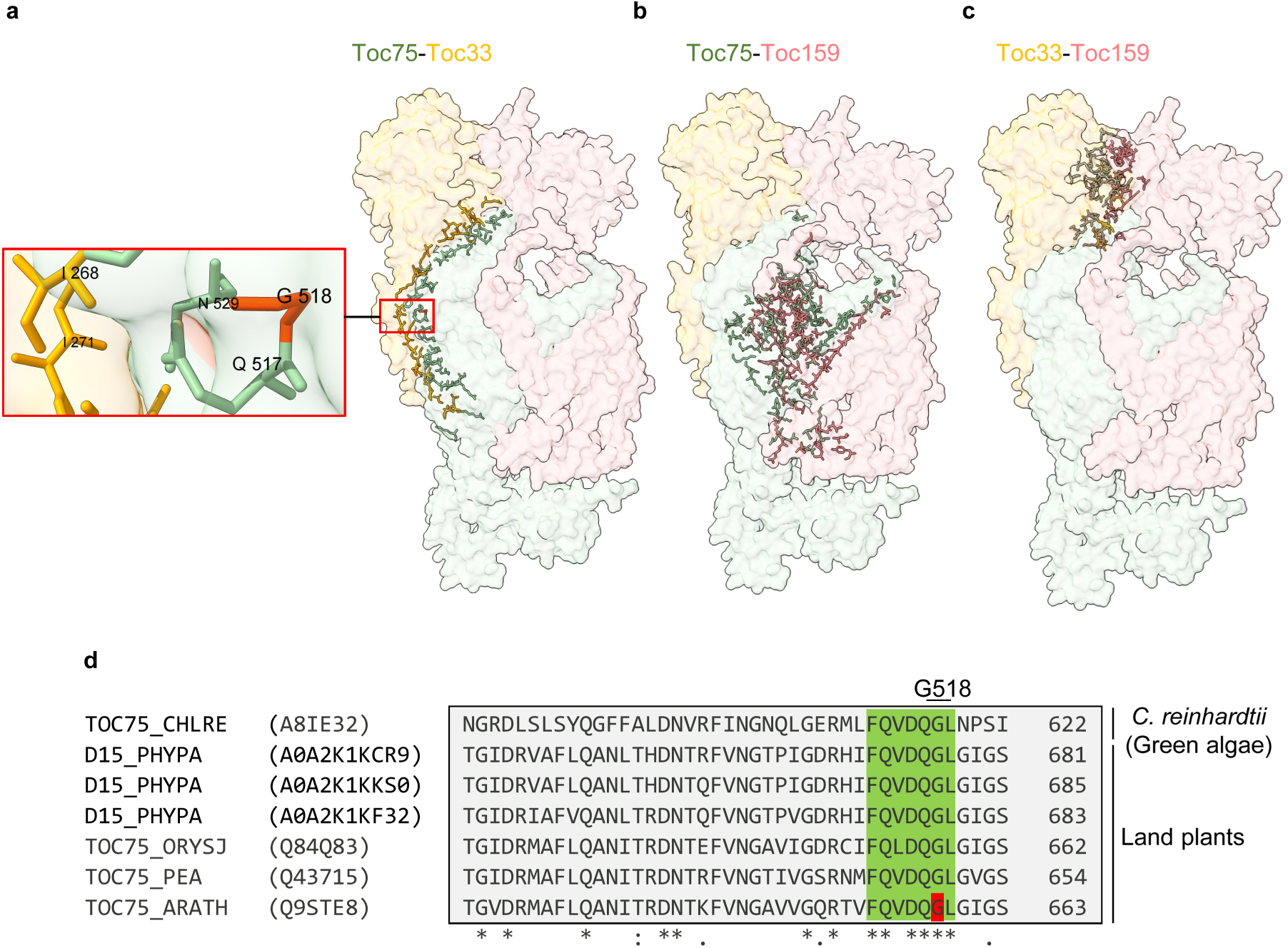
Toc75 acts as a central hub of interactions with Toc33 and Toc159. **a-c.** Surface models of the TOC-P complex (front view) that have been annotated to highlight residues participating in inter-subunit interactions. Interacting residues are shown in stick model format with darker colours. The models show that Toc75-Toc33 contacts occur in the membrane and cytosolic regions (**a**); that Toc75-Toc159 contacts are numerous in the membrane region (**b**); and that Toc33-Toc159 interactions are limited to the cytosolic region, between the GTPase domains (**c**). The inset in **a** shows the position of a Toc75 missense mutation present in the *kd-toc75* (*mar1/toc75-III-3*) mutant (G518R of mature Toc75; G658R of the precursor); the G518 residue is highlighted in dark orange/red. This mutation destabilizes the TOC-P and TOC-N complexes, as shown in Fig. 2. **d.** Multiple sequence alignment (MSA) of Toc75-related sequences showing conservation of glycine 518. Toc75-related sequences were obtained from Uniprot for *C. reinhardtii* (A8IE32; TOC75_CHLRE), *P. patens* (A0A2K1KCR9, D15_PHYPA; A0A2K1KKS0, D15_PHYPA; A0A2K1KF32, D15_PHYPA), *O. sativa* (Q84Q83, TOC75_ORYSJ), *P. sativum* (Q43715, TOC75_PEA), and *A. thaliana* (Q9STE8, TOC75-3_ARATH). Complete sequences were aligned in each case, but for simplicity only regions of interest are shown here. Clustal Omega was used to perform the MSA^54^, and the relevant regions were copied out manually. Coordinates at right show positions in each protein sequence; symbols at the bottom (*:.) indicate the degree of conservation at each position. It is evident that G518 (red) of *A. thaliana* Toc75 is highly conserved, and present in a conserved motif (green box). This is in accordance with the observation that G518 is important for stabilization of the TOC complexes (Fig. 2).

**Extended Data Fig. 7.**
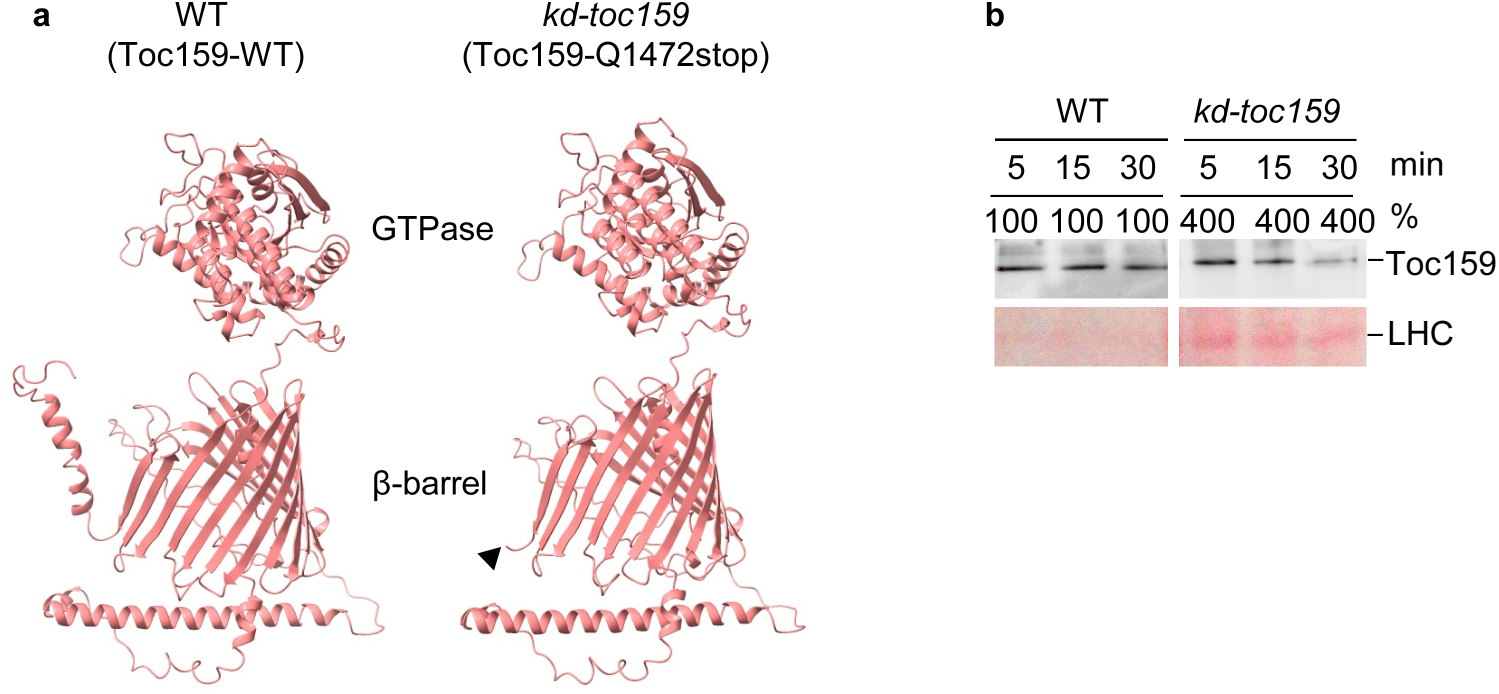
The C-terminal transmembrane helix of Toc159 is required for stabilization of Toc159 in the membrane. **a.** In the TOC-P structure, Toc159 is comprised of a cytosolic GTPase domain, a membrane-embedded β-barrel domain, and two additional alpha helices at the membrane: the first lies along the membrane facing towards the intermembrane space, and the second is a C-terminal transmembrane helix (TMH). The C-terminal TMH (32 residues in length) is absent in the *kd-toc159* (*fts1/ppi2-3*) mutant because of a mutation causing a premature stop codon (Q1472stop). The black triangle indicates the position of the truncation. **b.** Stability of the mutant Toc159 protein is substantially compromised. Membrane fractions from WT and *kd-toc159* plants were subjected to alkaline treatment (100 mM Na_2_CO_3_) for the indicated time periods, and then analysed by immunoblotting. The mutant protein was extracted much more readily than the WT protein, indicating instability. It was necessary to load four times more of the *kd-toc159* samples (see percentage values) in order to clearly detect Toc159 by immunoblotting; this indicated that the mutant protein is present at much lower levels than the WT protein, which may be related to its instability or reflect additional defects in biogenesis. Detection of LHC protein by Ponceau staining was used to normalize loading of the mutant samples.

**Extended Data Fig. 8.**
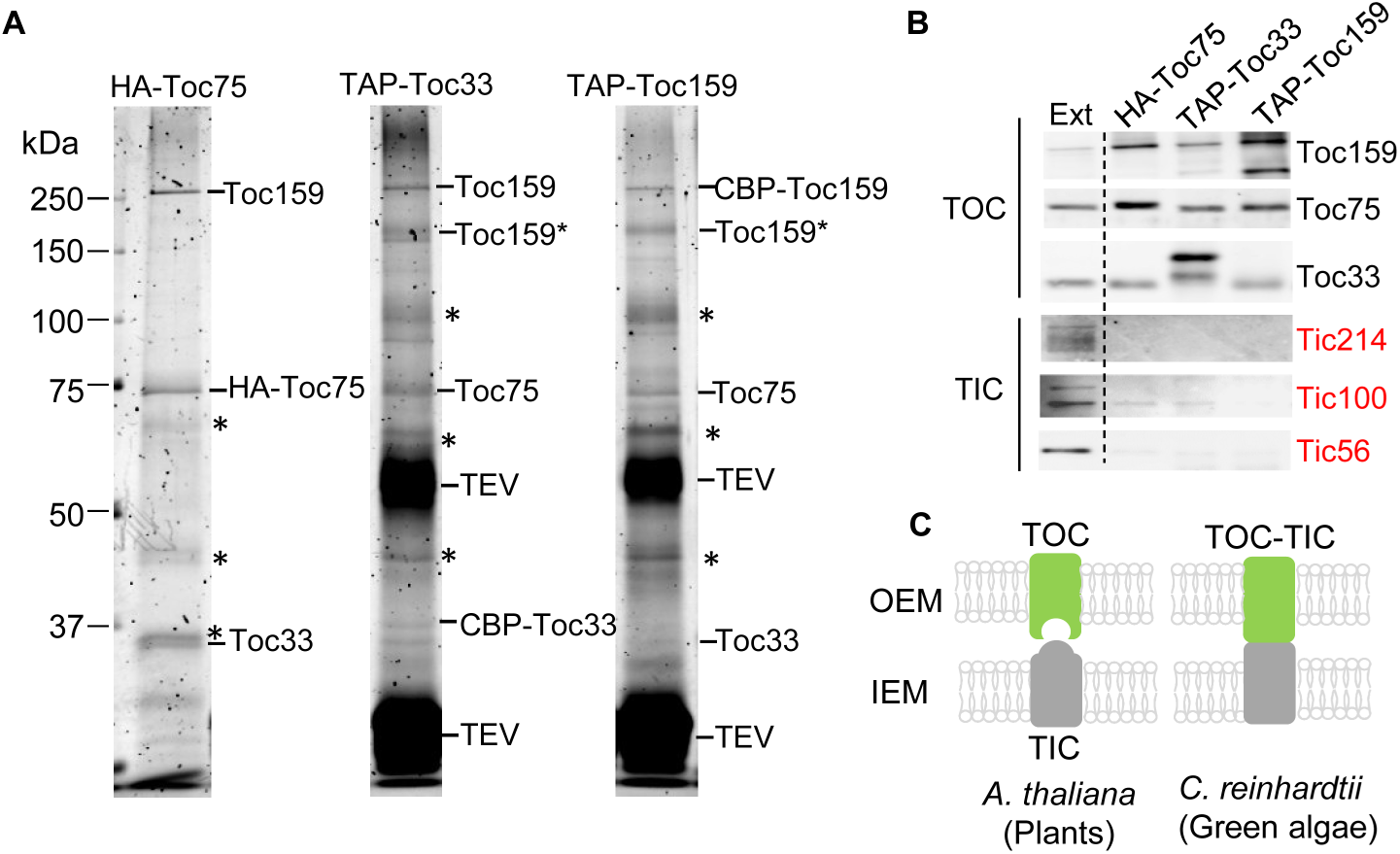
Analysis of TOC-TIC interactions in *A. thaliana* reveals that the two plant complexes are separable. **a.** Staining analysis of affinity-purified TOC complexes from three different transgenic lines. Chloroplast-enriched samples were prepared from 2-week-old transgenic plants expressing HA-tagged Toc75 or tandem-affinity-purification (TAP)-tagged Toc33 or Toc159. Samples were solubilized with 1% β-DM to extract membrane protein complexes before purification. The TAP tag carries two affinity domains (calmodulin-binding peptide (CBP), and the terminal immunoglobulin G (IgG)-binding Protein A domain) separated by a tobacco etch virus (TEV) protease cleavage site. Excess TEV protease was used to elute the TAP samples after Protein A-based purification, and the protease is clearly present in the relevant samples including in aggregated form. Resolved polypeptides were visualized by Flamingo fluorescent staining. Asterisks indicate non-specific interactions, whereas Toc159* indicates a common proteolytic fragment of Toc159. Positions of molecular weight markers (sizes in kDa) are indicated at left. Staining at the expected positions for prominent TIC components was not detected. **b.** Immunoblotting analysis of the affinity-purified TOC complex samples. Samples similar to those shown in **a** were analysed by immunoblotting using antibodies against TOC and TIC components. Whereas the TOC components were all clearly present as expected, the TIC components were either completely absent or present at negligible levels. Ext indicates a comparable extract from WT plants (not subjected to affinity purification) for comparison. **c.** Model comparing the relationship between TOC and TIC in plants and algae. Results in **a** and **b** indicated that the TOC and TIC complexes are readily separable in plants, and clearly are not deeply integrated and fused as they are in green algae.

**Extended Data Fig. 9.**
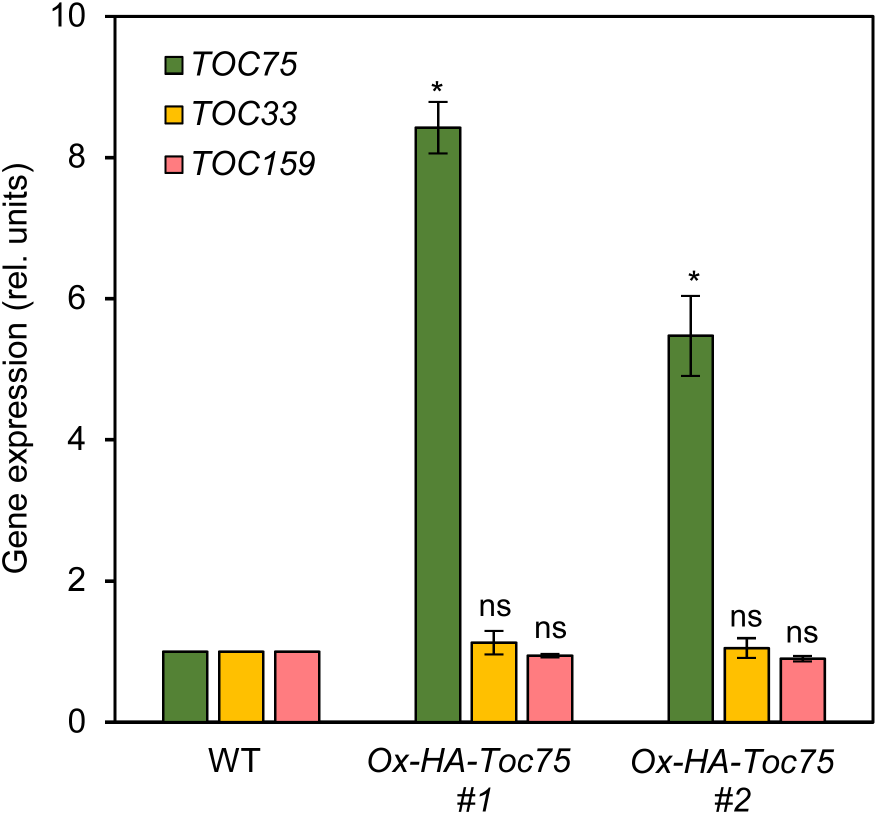
Analysis of *TOC* gene expression in the two *Ox-HA-Toc75* over-expressor lines. Gene expression analysis was performed by qRT-PCR, using RNA samples extracted from 2-week-old plants. The relevant transgenic plant lines were identified in a screen for complemented *HA-Toc75* lines, and are also presented in Fig. 1 (lines #2 and #3). Expression data for *TOC* genes were normalized using data for *ACTIN2*. Asterisks indicate significance according to paired one-tailed Student’s t tests comparing the transgenic lines with WT (\**p* < 0.01; ns, not significant). All values are means ± SEM (n = 3 experiments). The data show that *TOC* genes (apart from *TOC75*) are not generally over-expressed in the *Ox-HA-Toc75* lines.

**Extended Data Fig. 10.**
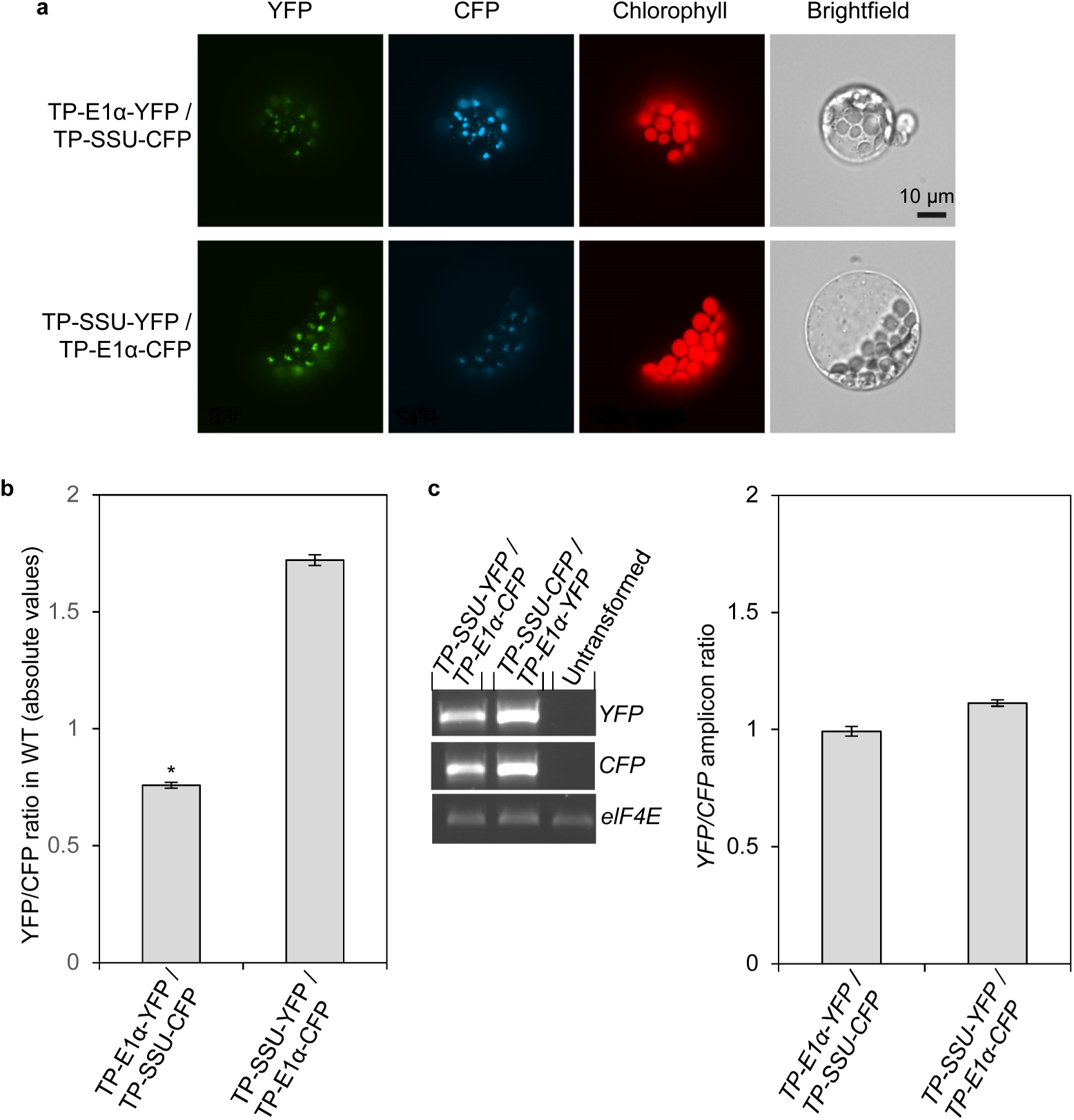
Ratiometric imaging analysis reveals that transit peptide identity influences chloroplast protein import efficiency in vivo. **a.** Qualitative analysis of the relative chloroplast-targeting efficiencies of the E1α and SSU transit peptides. Protoplasts from WT were co-transfected with plasmids encoding the indicated transit peptide (TP) fusions, and fluorescence microscopy was used to detect the transiently-expressed proteins. The YFP (green; left panels) and CFP (cyan; centre-left panels) fluorescence signals were recorded, alongside chlorophyll autofluorescence (red; centre-right panels) and brightfield images (right panels). Each set of images is of the same cell, and is representative of a large number of images derived from four independent experiments. **b.** Quantitative analysis of the experiments presented in **a**. The YFP/CFP fluorescence intensity ratios within individual chloroplasts were quantified using image analysis software; the data values shown here are absolute ratio values that have not been normalized. For each combination of constructs, the data shown comprise ∼1000-2000 individual ratio measurements, derived from a total of up to 59 different protoplasts from four separate experiments. Asterisks indicate significance according to two-tailed Student’s t tests comparing the TP-E1α-YFP/TP-SSU-CFP with TP-SSU-YFP/TP-E1α-CFP (\**p* < 0.001). All values are means ± SEM (n = 1201-1865). **c.** Analysis of the gene expression of the transfected plasmids in **a** and **b**. Samples of protoplasts that were either untransformed or had been co-transfected with the construct-pairs described above were subjected to RT-PCR analysis using construct-specific primers; similar analysis using *eIF4E1*-specific primers provided a positive control for the untransformed sample. Control PCR reactions containing the RNA samples as template (i.e., no reverse transcription) produced no amplification using the construct-specific primers. Typical stained agarose gels are shown at left, while quantitative analysis of the relevant amplicons in repeated experiments is shown at right. Corresponding values for *YFP* and *CFP* were used to calculate *YFP/CFP* ratios, which were then used to calculate means. Presented values are mean ratios ± SEM (n = 5 experiments). The data show that the two transfected genes (*YFP* and *CFP*) were expressed at a similar level in co-transfected cells of the same experiment.

## METHODS

### Plant materials and growth conditions

All plants used were *Arabidopsis thaliana* (Columbia-0). For most experiments, plants were grown for 7-14 days on Murashige and Skoog (MS) medium under long-day photoperiods (16-h light / 8-h dark; 150 µmol m^−2^ s^−1^) and at 22°C. For propagation, plants were grown on soil under similar conditions in a greenhouse. The following, previously-described TOC mutant genotypes were employed in this study: *ko-toc33* (*ppi1-1*)^14^, *ko-toc34* (*ppi3-2*)^12^, *kd-toc159* (*fts1*, *ppi2-3*)^20,24^, *kd-toc75* (*mar1*, *toc75-III-3*)^22,23^, *RNAi-toc75* (atToc75-III↓#6)^22^, and *ko-toc132-2* (*toc132-2*)^15^. The generation of the *HA-Toc75* and *Ox-HA-Toc75* lines is described below.

For de-etiolation experiments, seeds were germinated and grown in the dark at 22°C for 7 days as previously described^18^. Then, the dark-grown seedlings were exposed to light-dark cycles (16-h light / 8-h dark) for 7 days. Samples were collected at different time points to conduct gene expression analysis.

### Generation of *HA-Toc75* transgenic lines

Genomic DNA corresponding to the entire *TOC75* gene (At3g46740), including the native promotor and untranslated regions, was amplified by PCR from wild-type leaves using primers (gToc75-F/gToc75-R) designed to anneal 1005 bp upstream of *TOC75* start codon and 504 bp downstream of *TOC75* stop codon, and then cloned into a pGEM-T easy vector (Promega) **(Supplementary Table 2a)**. A nucleotide sequence (5’-TATCCTTATGATGTTCCTGATTATGCT-3’) encoding a haemagglutinin (HA)-peptide was inserted between codons 2 and 3 of the mature Toc75 protein (142E-HA-143E) by inverse PCR using a set of primers: mToc75-HA-F and mToc75-HA-R **(Supplementary Table 2a)**. Then, the full-length *HA-TOC75* sequence was cloned into the plant binary vector pPZP221, generating a transformation vector, pPZP221-HA-Toc75^14,55^. Following transformation into agrobacterium, the plants were transformed by floral dip method as reported earlier^14,56^. *HA-Toc75* transformants were selected on MS medium containing 110 μg mL^−1^ gentamycin, and homozygous T3 plants were obtained after segregation analysis. The *HA-Toc75* transgenic lines exhibited different levels of *TOC75* expression. The second and third lines exhibited unusually high levels of *TOC75* expression **(Extended Data Fig. 9)** and were therefore employed as over-expressor lines (*Ox-HA-Toc75*, #1 and #2).

### Affinity purification of HA-Toc75 and TAP-Toc33

To purify HA-Toc75/TAP-Toc33 complexes, a crude chloroplast preparation was made as described^57^. Seedlings were homogenized in chloroplast isolation (CI) buffer (0.3 M sorbitol, 5 mM magnesium chloride (MgCl_2_), 5 mM ethylene glycol tetraacetic acid (EGTA), 5 mM ethylenediaminetetraacetic acid (EDTA), 10 mM sodium hydrogen carbonate (NaHCO_3_), 20 mM 4-(2-hydroxyethyl)piperazine-1-ethanesulfonic acid (HEPES)-KOH, pH 8.0) and centrifuged at 1000×g for 5 min. The pellet was resuspended in HMS (HEPES-MgSO_4_-sorbitol (HMS) buffer (50 mM HEPES-NaOH, pH 8.0, 3 mM magnesium sulfate (MgSO_4_), 0.3 M sorbitol^57^. Alternatively, whole cell extracts were prepared from roots or etiolated seedlings using the CI and HMS buffers as described above^57^.

Then, the samples were solubilized with 1% (w/v) β-dodecyl maltoside (DDM) for 10 min and centrifuged at 20,000×g for 10 min^38^. Supernatant (500 µL) was loaded onto a small 1 mL spin-column containing 20-50 µL of anti-HA beads (HA-tagged protein purification kit [code: 3320], MBL) for HA-Toc-75 or IgG-sepharose beads (IgG-sepharose 6 fast flow [code, 17-0969-01], Cytiva) for TAP-Toc33, and the samples were mixed end-over-end for 1 h. Flow-through was removed by centrifugation for 10 s in a microcentrifuge. Then, the column was washed for 3-5 times with wash buffer (MBL). HA-Toc75 samples were eluted with an elution buffer containing HA peptide (MBL). TAP-Toc33 samples were eluted after digesting with TEV protease (7.5 U/μL) for 2 h at 16°C^20^. After elution, the samples were concentrated with 30 kDa cut-off ultrafilters (Sartorius). All the extraction and purification steps were performed at 4°C unless specified.

### Protein extraction, SDS-PAGE and immunoblotting

Protein samples were prepared from different plant tissues by homogenization in protein extraction buffer (50 mM Tris-HCl, pH 7.6, 1 mM phenylmethylsulfonyl fluoride (PMSF), 10% (v/v) glycerol, 0.1% (w/v) β-DM, 1× protease inhibitor cocktail (Sigma)). Polyacrylamide gels (10-15%) were casted using high-tris gel systems as reported previously^58^. Proteins were heat-denatured at 95°C in the presence of 100 mM dithiothreitol (DTT) and 2% (w/v) sodium dodecyl sulphate (SDS), for either 3 min (for purified samples) or 5 min (for whole cell extracts). After electrophoresis, proteins were visualized by staining with either Flamingo fluorescent dye (Bio-Rad), silver nitrate, or Coomassie brilliant blue. Alternatively, proteins were transferred onto nitrocellulose membrane to enable probing with different antibodies. The primary antibodies used in this study were as follows: anti-atToc75-III POTRA-domain (1:1000)^20,59^; anti-atToc159 A-domain (1:5000)^20,59^; anti-atToc33 peptide (1:500)^20,59^; atToc34 (Agrisera, AS07 238) (1:2000); anti-atToc132 A-domain (1:1000)^20,59^; anti-atToc120 A-domain (1:1000)^59^; anti-atTic110 stromal domain (1:5000)^59^; anti-atTic40 stromal domain (1:100000)^59^; anti-atTic56 (1:5000)^60^, anti-atTic100 (1:5000)^60^, anti-atTic214 (1:5000)^60^, anti-actin (1:3000; AS132640, Agrisera); anti-histone H3 (1:1000; ab1791, Abcam); and anti-HA (1:1000; H6908, Sigma). The secondary antibody was anti-rabbit IgG conjugated with horseradish peroxidase (Sigma, 12-348) (1:5000). Immunodetection was by chemiluminescence and monitored using an ImageQuant LAS-4000 imager (GE Healthcare). The protein bands (staining/radiolabelling) were quantified using ImageJ^61^ and the raw data was exported to excel and analysed **(Figs. 2a,b, 4b-e and 5c-e)**.

Membrane fractions from wild-type and *kd-toc159* mutant leaf extracts were prepared by as described. These fractions were washed with 100 mM Na_2_CO_3_ for up to 30 min. Samples were centrifuged (22,000×g) for 30 min, and the pelleted fractions were analysed by SDS-PAGE and immunoblotting as described above.

### Gene expression analysis by qRT-PCR

RNA extractions were performed using a Spectrum Plant Total RNA kit (Merck). Reverse transcription was performed by using SuperScript IV reverse transcriptase (Invitrogen). For quantitative reverse transcriptase PCR (qRT-PCR), a qPCRBIO SyGreen Mix Hi-ROX kit (PCR Biosystems Ltd.) was employed, together with a StepOnePlus Real-Time PCR System (Applied Biosystems) and its associated software according the manufacturer’s instructions. The primers used for PCR amplification are shown **(Supplementary Table 2b)**. Expression data for genes of interest were normalized using data for *ACTIN2* (At3g18780).

### Radiolabelling

For each genotype, forty 6-7-day-old seedlings were collected into 3 mL MS liquid medium in a 6-well microtitre plate. Seedlings were incubated with 30 µCi/mL of Na_2_^35^SO_4_ for 5-16 h at

22°C under 80-100 µmol photons m^−2^ s^−1^ white light^38,62^. Radiolabelling was stopped by adding 100 mM Na_2_SO_4_. Seedlings were collected into a 1.5 mL microfuge tube and homogenized using a small plastic pestle. Protein extraction was performed as described earlier, and the seedling extract was further subjected to anti-HA affinity purification also as described earlier. After affinity purification, the samples were resolved by SDS-PAGE and then imaged by staining or phosphorimaging (Storm Molecular Imager, Molecular Dynamics).

### Protoplast isolation, transfection and analysis

The transit peptide fusions of the Rubisco small subunit (SSU; At1g67090) and pyruvate dehydrogenase E1α subunit (E1α; At1g01090) to YFP and CFP were generated using the p2GWY7^63^ and p2GWC7 vectors^63^ by Gateway cloning **(Supplementary Table 2c)**. In each case, at least seven residues of the mature part of the preprotein were retained, so as not to disrupt the stromal processing peptidase cleavage site or targeting efficiency. The transit peptide coding sequences were amplified from first-strand cDNA or plasmid cDNA clones using gene-specific primers **(Supplementary Table 2c)**. pI values of the TPs of SSU and E1α in **Fig. 7** were calculated using the ExPASy online server (https://web.expasy.org/compute_pi/)^64^.

Protoplasts were prepared from 2-4-week-old plants and transfected using well established methodologies^65^. Transfections employed 5 µg plasmid DNA (per construct) and 100 µL protoplast suspension (2 × 10^5^ cells). The transformed protoplasts were analysed by either immunoblotting, RT-PCR, or fluorescence microscopy.

For immunoblotting, the protoplasts were suspended in protein extraction buffer (100 mM Tris pH 6.8, 10% (v/v) glycerol, 0.5% (w/v) SDS, 0.1% (v/v) Triton X-100, fresh 10 mM DTT, fresh 1× protease inhibitor cocktail (Sigma)) and vortexed vigorously for 1 min. The lysates were denatured, before subjecting them to SDS-PAGE and immunoblotting as described earlier.

For RT-PCR analysis of transient expression, total RNA was isolated from transfected (or untransformed) protoplasts using an RNeasy Plant Mini kit according to the manufacturer’s instructions (Qiagen). DNAse І treatment and first-strand cDNA synthesis (using 3 μg RNA) were conducted as described previously^66^. To assess expression of the full-length fusions, RT-PCR employed primers specific for YFP as follows **(Supplementary Table 2c,d)**: YFP-F/YFP-R. To assess expression of the target peptide fusions, RT-PCR employed primers specific for each construct as follows: SSU-F, E1α-F, and CFP/YFP-R. In each case, the *eIF4E1* (At4g18040) gene was employed as a housekeeping control for normalization purposes, using the following primers: eIF4E1-F/eIF4E1-R. Following electrophoresis, bands were visualized by staining with SYBR Safe (Invitrogen) and quantified using Aida software (Raytest).

For fluorescence microscopy, samples were analysed 16-24 hours after co-transfection using a Nikon Eclipse TE-2000E inverted fluorescence microscope equipped with filters for analyzing YFP (exciter HQ500/20x, emitter HQ535/30m), CFP (exciter D436/20x, emitter D480/40m), and chlorophyll autofluorescence (exciter D480/30x, emitter D660/50m) (Chroma Technologies). For each plasmid combination, image sets of co-expressing protoplasts were captured in at least three independent transfection experiments. The ratio of exposure times for YFP and CFP was kept at 1:1 throughout all experiments and for all genotypes. Exposure times ranged from 100 ms to 500 ms, depending on brightness of signal, and the gain setting was zero at all times; the YFP images were always captured first. Quantification of the YFP:CFP signal intensity ratios was conducted using Openlab software (Improvision). Firstly, ratio images of corresponding YFP/CFP images pairs were generated using the “Ratio” software module; all ratios were measured as YFP over CFP. To avoid unspecific skewing of the ratio data for individual chloroplasts, the “Density Slice” module was employed, which allows the specific selection of signal spots within chloroplasts (this was necessary because we observed that the fluorescence within chloroplasts was not uniformly distributed). A density slice layer was generated for each image pair, and then applied for the extraction of the ratio data from individual signal spots. Cells expressing only one of the two fluorophores were rare (∼10%), and were not analysed.

### Negative staining by electron microscopy (EM)

A thin clean film of carbon on top of a copper mesh 200 (C101/100, TAAB) was glow discharged for 25 s at 15 mA current (Pelco, easiGlow) before 3 μL of a freshly-purified TOC-P sample was applied directly onto the grid. After 1 min of incubation at ambient temperature, the sample was absorbed by touching the side of the grid with a wedge of filter paper (Whatman, 1001-090). The grid was rapidly dropped, face downwards, on top of a 20 μL drop of 2% uranyl acetate solution dispensed onto the surface of Parafilm, where it was then incubated for 10 s. The excess stain was blotted away until a thin film remained, and the grid was then air-dried for 5 min. A total of 195 images were collected at a nominal magnification of 50,000 and a pixel size of 2.3 Å on a 200 kV electron microscope (JEM-2100Plus).

### EM data processing

The raw images were processed in CryoSPARC^67^, where CTFFIND4^68^ was used for Contrast Transfer Function (CTF) estimation using a higher amplitude contrast of 0.25 at a minimum resolution of 50 Å . Initially, particles were either picked manually or using the blob picker with a particle diameter in the range of 50-200 Å. The best classes were then selected for the template picker using 150 Å particle diameter. After extracting about 20,000 particles with a box size of 200 pixels, two rounds of 2D classifications were run with a constant CTF for output particles. The best 2D classes comprising about 8,000 particles were used for comparison with 2D projections of the AF3-generated model of the TOC-P complex.

### Crosslinking mass spectrometry

The HA-purified TOC complex was crosslinked with 3 mM bis(sulfosuccinimidyl)suberate (BS3) (Thermo Scientific) at room temperature for 3 h. The reaction was terminated by adding 50 mM Tris. Detergent (DDM) was removed by using HiPPR detergent removal resin (Thermo Scientific) prior to the preparation of the samples for mass spectrometry (MS). The crosslinked samples were denatured in 4 M urea (in 100 mM ammonium bicarbonate, pH 7.8) for 10 min, before cysteine reduction with 10 mM Tris(2-carboxyethyl)phosphine (TCEP) for 30 min at room temperature, and alkylation with 50 mM 2-chloroacetamide for 30 min at room temperature in the dark. Next, the samples were pre-digested with endoproteinase LysC (Wako) (1:100 µg enzyme:protein ratio) at 37°C for 2 h with shaking and then further digested with trypsin (Promega) (1:40 µg enzyme:protein ratio) for 16 h with shaking. The digested peptides were loaded onto C18 stage tips, pre-activated with 100% acetonitrile and 0.1% trifluoroacetic acid. After centrifugation, peptides were washed with 0.1% trifluoroacetic acid and eluted in 50% acetonitrile / 0.1% trifluoroacetic acid. Eluted peptides were dried in a speed-vac. Dried peptides were resuspended into 5% acetonitrile / 5% formic acid before LC-MS/MS analysis.

Peptides were separated by nano liquid chromatography (Thermo Scientific Ultimate RSLC 3000) coupled in-line to a QExactive mass spectrometer equipped with an Easy-Spray source (Thermo Fischer Scientific). Peptides were trapped onto a C18 PepMac100 precolumn (300 µm i.d., 5 mm length, 100Å, Thermo Fisher Scientific) using Solvent A (0.1% formic acid, HPLC grade water). The peptides were further separated onto an Easy-Spray RSLC C18 column (75 µm i.d., 50 cm length, Thermo Fisher Scientific) using 60 and 120 min linear gradients (15% to 35% Solvent B (0.1% formic acid in acetonitrile)) at a flow rate 200 nL/min. The raw data were acquired on the mass spectrometer in a data-dependent acquisition (DDA) mode. Full-scan MS spectra were acquired in the Orbitrap (scan range 350-1500 m/z, resolution 70,000, AGC target 3e6, maximum injection time 50 ms). The 10 (60 min gradient) and 20 (120 min gradient) most intense peaks were selected for higher-energy collision dissociation (HCD) fragmentation at 30% of normalized collision energy. HCD spectra were acquired in the Orbitrap at resolution 17,500, AGC target 5e4, maximum injection time 120 ms with fixed mass at 180 m/z. Charge exclusion was selected for unassigned and 1+ ions. The dynamic exclusion was set to 20 s and 40 s for 60 min and 120 min gradients, respectively.

Acquired tandem mass (MS/MS) spectra were searched using pLINK3^69^ and MetaMorpheus 1.0.9^70^, against an *Arabidopsis thaliana* database (Proteome ID UP000006548) downloaded from UniProt (2024_03_14), containing 18,300 protein entries and the tagged Toc75 protein sequence. In addition, 292 contaminant sequences were added to the database while processing with pLINK3. The default contaminant database of MetaMorpheus was used. The following variable modifications were set: Oxidation (M), Acetylation (N-ter). Carbamidomethylation was set as a fixed modification. Data were filtered at FDR below 5%. Trypsin was set as the endoprotease. All identified crosslinked spectra were manually inspected and validated. The TOC crosslinked spectra are displayed in **Supplementary Fig. 8**.

### Alphafold3 analysis

AF3 predictions were performed in the on-line server (https://alphafoldserver.com)^25^. To generate TOC-P models, the component sequences of TOC-P were submitted in the following order; 1. the mature protein sequence of Toc75 (D141-Y818); 2, truncated Toc159 (E781-Y1503); 3, full-length Toc33 (M1-297L); and two GTP ligands **(Fig. 3a)**. To generate TOC-N models, the component sequences of TOC-N were submitted in the following order **(Extended Data Fig. 2)**; 1. the mature protein sequence of Toc75 (D141-Y818); 2, truncated Toc132 (A486-Q1205); 3, full-length Toc34 (M1-S313); and two GTP ligands. TOC complex protein sequences from green alga, *C. reinhardtii*, were also submitted in the following order; 1, the mature protein sequence of Toc75 (E128-F798); 2, truncated Toc90 (P190-L967); 3, full-length Toc34 (M1-D396), and two GTP ligands **(Fig. 3e)**. For each predictions, five models were generated and ranked in order of their scores. Based on their ranking scores and architecture, they were found be similar overall. The top-scored models were thoroughly studied to reveal the structural features by Chimera-1.17 (https://www.cgl.ucsf.edu/chimera/). The models show in the manuscript are all derived from Top-scored models. Surface electrostatic potential was calculated using Chimera-1.17.

### Phyre2 and I-TASSER analysis

The membrane domain of Toc159 (residues 1045-1503) was submitted to the Phyre2 protein threading server (http://www.sbg.bio.ic.ac.uk/phyre2/html/page.cgi?id=index)^52^ and to the I-TASSER protein structure and function prediction server (https://zhanggroup.org/I-TASSER/)^53^. The models generated by the Phyre2 server were analysed by Chimera as above.

### Molecular dynamics simulations

The AF3 TOC-P model shown in **Fig. 3c** was shortly minimized with position restraints of 1000 kJ mol^−1^ nm^−2^, with the CHARMM36 forcefield converging after 118 steps. The resulting structure was used as a starting point for the coarse-grained (CG) modelling. The topologies to run CG simulations were generated with the martinize tool choosing the Martini v.2.2 force field with an elastic network^71,72^. The force bond constant was set to 500 kJ mol^−1^ nm^−2^ with lower and upper elastic bond cut-offs of 0.5 and 0.9 nm, respectively. Elastic bonds were checked to make sure linker regions were kept flexible. The protein was then embedded in a symmetric POPC bilayer using the insane tool and solvated in a 150 mM NaCl solution resulting in a box of dimensions 20 × 20 × 20 nm^3^ **(Fig. 3h)**. Minimization was performed for 1000 steps with steepest-descent and position restraints of 1000 kJ mol^−1^ nm^−2^, followed by a step-wise equilibration increasing the time-step from 10 fs to 20 fs and decreasing position restraints. During the equilibration process, the pressure was semi-isotropically coupled via a Berendsen barostat to a reference value of 1 bar with a coupling constant of 5 and compressibility of 3e^−4^ bar^−1^, while the temperature was coupled in the same way as the production run. Finally, production run was performed at 323 K and 1 bar. Temperature was maintained via V-rescale thermostat with a coupling constant of tau_t = 1 ps, whereas semi-isotropic pressure coupling was achieved via a Parinello-Raham barostat with a coupling constant tau_p = 12 ps and compressibility 3e-4 bar^−1^. Three replicates of coarse-grained molecular dynamics simulations were run in GROMACS v2021.4^73^ for 10 µs each.

Root mean square deviations were calculated in GROMACS using the gmx rms tool considering only backbone beads. The RMSD values of the GTPase domains of interest were computed and plotted after fitting on the whole transmembrane region of the modelled TOC complex. For this analysis, the linker regions were defined to include only unstructured elements of the proteins spanning the following residues: linker-1 of Toc33, D251-S260; linker-2 of Toc159, L1077-R1086 (amino acid coordinates in the AF3 truncated Toc159 structure are L297-R306). Lipid-protein contact analysis was computed over the whole 10 µs-long simulation **(Supplementary Videos 1 and 2)**. A contact was defined when any protein CG bead was within a distance of up to 0.6 nm from any POPC bead. The total contact counts were divided by the number of frames to report an averaged value per frame.

VMD^74^ was used for visualization of all systems. Ad hoc Python 3 scripts (containing MDAnalysis^75^ and NumPy libraries) were used during post-processing and data analysis. Matplotlib^76^ was used for plotting.

### Sequence retrieval and analysis

Gene sequences for the following proteins from *A. thaliana* were used in this study for experimental or computational analysis: Toc75 (At3g46740), Toc159 (At4g02510), Toc33 (At1g02280), Toc34 (At5g05000), Toc120 (At3g16620), Toc132 (At1g16640), Tic214 (AtCg01130), Tic100 (At5g22640), Tic56 (At5g01590), Tic110 (At1g06950), Tic40 (At1g16620), LHCB1.1 (At1g29920), Actin2 (At3g18780), Rubisco small subunit SSU (At1g67090), and pyruvate dehydrogenase E1α subunit (At1g01090). Gene sequences for the following proteins from *C. reinhardtii* were used in this study for computational analysis: Toc75 (Cre03.g175200), Toc90 (Cre17.g734300.t1.1), and Toc34 (Cre06.g252200).

Sequences were obtained from the TAIR (http://www.arabidopsis.org/), Phytozome (https://phytozome.jgi.doe.gov/pz/portal.html), Uniprot (https://www.uniprot.org/), or National Center for Biotechnology Information (NCBI) (https://www.ncbi.nlm.nih.gov/) databases.

Multiple sequence alignments were performed using Clustal Omega (https://www.ebi.ac.uk/jdispatcher/msa/clustalo) using default settings^54^.

## Acknowledgements

This research was supported by UK Research and Innovation (UKRI) Biotechnology and Biological Sciences Research Council (BBSRC; grants B/F020325/1, BB/K018442/1, BB/N006372/1, BB/R009333/1, BB/R016984/1, BB/V007300/1, BB/W015021/1, BB/X000192/1 and BB/Z516624/1 to R.P.J.). In addition, S.N. was supported by a Marie-Curie Individual Fellowship (TOC maker, 894893) from the European Commission, a John Fell Fund award (APD00260) from the University of Oxford, and a JSPS KAKENHI grant (16K21737) from Japan; and P.P. was supported by the Spanish Agencia Estatal de Investigacion (CNS2022-135817). S Khalid’s group was supported by UKRI Engineering and Physical Sciences Research Council (EPSRC; grants EP/X035603, EP/V030779/1 and EP/Y008693/1) and the Wellcome Trust. Access to High Performance Supercomputing was provided by HECBioSim through EPSRC (grant EP/X035603). I.A.’s group was supported by the Wellcome Trust (grants 223107 and 302632), BBSRC (grants BB/R007195/1 and BB/W016613/1), and Cancer Research United Kingdom (grant C35050/A22284). We thank Prof. Felix Kessler (Université de Neuchâtel, Switzerland) for the *TAP-Toc159* transgenic line; Dr. Jaypal Darbar (University of Oxford, UK) for assistance with plant growth and chloroplast preparations; Dr. Haoge Li (Shenyang Agricultural University, China) for assistance with the petal protein extractions; Dr. Yi Sun (University of Oxford, UK) for assistance with initiation of the TAP analyses; and Dr. Feijie Wu (formerly of University of Leicester, UK) for assistance with RT-PCR analyses of protoplast samples.

## Data availability

All data generated or analysed during this study are included in this published article or its supplementary information, apart from the following. The mass spectrometry proteomics data have been deposited to the ProteomeXchange Consortium via the PRIDE partner repository^77^ with the data set identifier PXD064453.

All unique materials are readily available from the authors.

## Author contributions

S.N. and R.P.J. conceived and designed the experiments. S.N. performed most of the experiments with assistance from the co-authors. S.N. and D.B. performed the AF3 analyses.

D.B. performed negative staining and EM data processing. S. Khalid conceived and supervised the MD simulations, while A.F.B. ran the simulations, analysed the data, and wrote the MD section. S. Kubis performed the ratiometric in vivo protein import assays using fusion protein pairs. Z.S. performed the gene expression analysis by quantitative qRT-PCR. S.C. performed immunoblotting analysis of the *RNAi-Toc75* and *Ox-HA-Toc75* transgenic lines. S. Kodru and U.F.P. assisted with the TAP and HA-tag affinity purifications. J.F. assisted with radioisotope labelling and HA-affinity purifications. P.P. performed segregation analysis of the *HA-Toc75* transgenic lines. V.R. and M.R performed the LC-MS/MS analysis. I.A. provided additional supervision of the AF3 analysis. R.P.J was responsible for overall supervision of the project. S.N and R.P.J. analysed the data and wrote the manuscript, and all authors approved it.

## Rights retention statement

This research was funded in whole or in part by UKRI (BB/F020325/1, BB/K018442/1, BB/N006372/1, BB/R009333/1, BB/R016984/1, BB/V007300/1, BB/W015021/1, BB/X000192/1 and BB/Z516624/1). For the purpose of Open Access, the author has applied a CC BY public copyright licence to any Author Accepted Manuscript (AAM) version arising from this submission.

## References

1. Jarvis, P. & López-Juez, E. Biogenesis and homeostasis of chloroplasts and other plastids. Nat. Rev. Mol. Cell Biol. 14, 787–802 (2013).

2. Sierra, J., Escobar-Tovar, L. & Leon, P. Plastids: diving into their diversity, their functions, and their role in plant development. J. Exp. Bot. 74, 2508–2526 (2023).

3. Kessler, F. & Schnell, D.J. The function and diversity of plastid protein import pathways: A multilane GTPase highway into plastids. Traffic 7, 248–257 (2006).

4. Li, H.M. & Chiu, C.C. Protein Transport into Chloroplasts. Annu. Rev. Plant Biol. 61, 157–180 (2010).

5. Nellaepalli, S., Lau, A.S. & Jarvis, R.P. Chloroplast protein translocation pathways and ubiquitin-dependent regulation at a glance. J. Cell Sci. 136(2023).

6. Shi, L.X. & Theg, S.M. The chloroplast protein import system: from algae to trees. Biochim. Biophys. Acta 1833, 314–31 (2013).

7. O’Neil, P.K. et al. The POTRA domains of Toc75 exhibit chaperone-like function to facilitate import into chloroplasts. Proc. Natl. Acad. Sci. USA 114, E4868–E4876 (2017).

8. Demarsy, E., Lakshmanan, A.M. & Kessler, F. Border control: selectivity of chloroplast protein import and regulation at the TOC-complex. Front. Plant Sci. 5(2014).

9. Jarvis, P. Targeting of nucleus-encoded proteins to chloroplasts in plants. New Phytol. 179, 257–285 (2008).

10. Sun, Y. & Jarvis, R.P. Chloroplast Proteostasis: Import, Sorting, Ubiquitination, and Proteolysis. Annu. Rev. Plant Biol. 74, 259–283 (2023).

11. Bauer, J. et al. The major protein import receptor of plastids is essential for chloroplast biogenesis. Nature 403, 203–207 (2000).

12. Constan, D., Patel, R., Keegstra, K. & Jarvis, P. An outer envelope membrane component of the plastid protein import apparatus plays an essential role in. Plant J. 38, 93–106 (2004).

13. Ivanova, Y., Smith, M.D., Chen, K.H. & Schnell, D.J. Members of the Toc159 import receptor family represent distinct pathways for protein targeting to plastids. Mol. Biol. Cell 15, 3379–3392 (2004).

14. Jarvis, P. et al. An Arabidopsis mutant defective in the plastid general protein import apparatus.Science 282, 100–103 (1998).

15. Kubis, S. et al. Functional specialization amongst the Arabidopsis Toc159 family of chloroplast protein import receptors. Plant Cell 16, 2059–2077 (2004).

16. Richardson, L.G.L. et al. Targeting and assembly of components of the TOC protein import complex at the chloroplast outer envelope membrane. Front. Plant Sci. 5(2014).

17. Fish, M. et al. New Insights into the Chloroplast Outer Membrane Proteome and Associated Targeting Pathways. Int. J. Mol. Sci. 23(2022).

18. Ling, Q.H., Huang, W.H., Baldwin, A. & Jarvis, P. Chloroplast Biogenesis Is Regulated by Direct Action of the Ubiquitin-Proteasome System. Science 338, 655–659 (2012).

19. Nellaepalli, S., Kim, R.G., Grossman, A.R. & Takahashi, Y. Interplay of four auxiliary factors is required for the assembly of photosystem I reaction center subcomplex. Plant J. 106, 1075–1086 (2021).

20. Ling, Q.H. et al. Ubiquitin-dependent chloroplast-associated protein degradation in plants. Science 363, 836 (2019).

21. Agne, B. et al. The Acidic A-Domain of Arabidopsis Toc159 Occurs as a Hyperphosphorylated Protein. Plant Physiol. 153, 1016–1030 (2010).

22. Huang, W.H., Ling, Q.H., Bédard, J., Lilley, K. & Jarvis, P. In Vivo Analyses of the Roles of Essential Omp85-Related Proteins in the Chloroplast Outer Envelope Membrane. Plant Physiol. 157, 147–159 (2011).

23. Stanga, J.P., Boonsirichai, K., Sedbrook, J.C., Otegui, M.S. & Masson, P.H. A Role for the TOC Complex in Arabidopsis Root Gravitropism. Plant Physiol. 149, 1896–1905 (2009).

24. Woodson, J.D. et al. Ubiquitin facilitates a quality-control pathway that removes damaged chloroplasts. Science 350, 450–454 (2015).

25. Abramson, J. et al. Accurate structure prediction of biomolecular interactions with AlphaFold 3. Nature 630(2024).

26. Richardson, L.G., Jelokhani-Niaraki, M. & Smith, M.D. The acidic domains of the Toc159 chloroplast preprotein receptor family are intrinsically disordered protein domains. BMC Biochem. 10, 35 (2009).

27. Jin, Z.Y. et al. Structure of a TOC-TIC supercomplex spanning two chloroplast envelope membranes. Cell 185, 4788–4800 (2022).

28. Liu, H., Li, A., Rochaix, J.D. & Liu, Z. Architecture of chloroplast TOC-TIC translocon supercomplex. Nature 615, 349–357 (2023).

29. Koenig, P. et al. The GTPase cycle of the chloroplast import receptors Toc33/Toc34:: Implications from monomeric and dimeric structures. Structure 16, 585–596 (2008).

30. Sun, Y.J. et al. Crystal structure of pea Toc34, a novel GTPase of the chloroplast protein translocon. Nat. Struct. Biol. 9, 95–100 (2002).

31. Weibel, P., Hiltbrunner, A., Brand, L. & Kessler, F. Dimerization of Toc-GTPases at the chloroplast protein import machinery. J. Biol Chem. 278, 37321–37329 (2003).

32. Shen, C. et al. Structural basis of BAM-mediated outer membrane beta-barrel protein assembly. Nature 617, 185–193 (2023).

33. Ganesan, I., Shi, L.X., Labs, M. & Theg, S.M. Evaluating the Functional Pore Size of Chloroplast TOC and TIC Protein Translocons: Import of Folded Proteins. Plant Cell 30, 2161–2173 (2018).

34. Ganesan, I. & Theg, S.M. Structural considerations of folded protein import through the chloroplast TOC/TIC translocons. Febs Lett. 593, 565–572 (2019).

35. Pipitone, R. et al. A multifaceted analysis reveals two distinct phases of chloroplast biogenesis during de-etiolation in *Arabidopsis*. elife 10(2021).

36. Solymosi, K. & Schoefs, B. Etioplast and etio-chloroplast formation under natural conditions: the dark side of chlorophyll biosynthesis in angiosperms. Photosynth. Res. 105, 143–166 (2010).

37. Weier, T.E. & Brown, D.L. Formation of the prolamellar body in 8-day, dark-grown seedlings. Am. J. Bot 57, 267–275 (1970).

38. Nellaepalli, S., Ozawal, S.I., Kuroda, H. & Takahashi, Y. The photosystem I assembly apparatus consisting of Ycf3-Y3IP1 and Ycf4 modules. Nat. Commun. 9(2018).

39. Lee, D.W., Lee, S., Oh, Y.J. & Hwang, I. Multiple Sequence Motifs in the Rubisco Small Subunit Transit Peptide Independently Contribute to Toc159-Dependent Import of Proteins into Chloroplasts. Plant Physiol. 151, 129–141 (2009).

40. Kim, J., Na, Y.J., Park, S.J., Baek, S.H. & Kim, D.H. Biogenesis of chloroplast outer envelope membrane proteins. Plant Cell Rep. 38, 783–792 (2019).

41. Wild, K. et al. Structural Basis for Conserved Regulation and Adaptation of the Signal Recognition Particle Targeting Complex. J. Mol. Biol. 428, 2880–97 (2016).

42. Liang K, J.Z., Zhan X, Li Y, Xu Q, Xie Y, Yang Y, Wang S, Wu J, Yan Z Structural insights into the chloroplast protein import in land plants. Cell 187, 1–14 (2024).

43. Ling, Q. et al. The chloroplast-associated protein degradation pathway controls chromoplast development and fruit ripening in tomato. Nat. Plants 7, 655–666 (2021).

44. Wallas, T.R., Smith, M.D., Sanchez-Nieto, S. & Schnell, D.J. The roles of toc34 and toc75 in targeting the toc159 preprotein receptor to chloroplasts. J. Biol. Chem. 278, 44289–97 (2003).

45. Smith, M.D., Hiltbrunner, A., Kessler, F. & Schnell, D.J. The targeting of the atToc159 preprotein receptor to the chloroplast outer membrane is mediated by its GTPase domain and is regulated by GTP. J. Cell Biol. 159, 833–843 (2002).

46. Hiltbrunner, A. et al. Targeting of an abundant cytosolic form of the protein import receptor at Toc159 to the outer chloroplast membrane. J. Cell Biol. 154, 309–316 (2001).

47. Tu, S.L. et al. Import pathways of chloroplast interior proteins and the outer-membrane protein OEP14 converge at Toc75. Plant Cell 16, 2078–2088 (2004).

48. Doyle, M.T. & Bernstein, H.D. Function of the Omp85 Superfamily of Outer Membrane Protein Assembly Factors and Polypeptide Transporters. Annu. Rev. Microbiol. 76, 259–279 (2022).

49. Reumann, S. & Keegstra, K. The endosymbiotic origin of the protein import machinery of chloroplastic envelope membranes. Trends in Plant Sci. 4, 302–307 (1999).

50. Reumann, S., Inoue, K. & Keegstra, K. Evolution of the general protein import pathway of plastids (Review). Mol. Membr. Biol. 22, 73–U20 (2005).

51. Bay, D.C., Hafez, M., Young, M.J. & Court, D.A. Phylogenetic and coevolutionary analysis of the β-barrel protein family comprised of mitochondrial porin (VDAC) and Tom40. Biochim. Biophys. Acta 1818, 1502–1519 (2012).

52. Kelley, L.A., Mezulis, S., Yates, C.M., Wass, M.N. & Sternberg, M.J. The Phyre2 web portal for protein modeling, prediction and analysis. Nat. Protoc. 10, 845–58 (2015).

53. Yang, J. et al. The I-TASSER Suite: protein structure and function prediction. Nat. Methods 12, 7–8 (2015).

54. Sievers, F. et al. Fast, scalable generation of high-quality protein multiple sequence alignments using Clustal Omega. Mol. Syst. Biol. 7(2011).

55. Hajdukiewicz, P., Svab, Z. & Maliga, P. The Small, Versatile Ppzp Family of Agrobacterium Binary Vectors for Plant Transformation. Plant Mol. Biol. 25, 989–994 (1994).

56. Clough, S.J. & Bent, A.F. Floral dip: a simplified method for Agrobacterium-mediated transformation of Arabidopsis thaliana. Plant J. 16, 735–43 (1998).

57. Flores-Pérez, U. & Jarvis, P. Isolation and Suborganellar Fractionation of Arabidopsis Chloroplasts. Methods Mol. Biol. 1511, 45–60 (2017).

58. Fling, S.P. & Gregerson, D.S. Peptide and Protein Molecular-Weight Determination by Electrophoresis Using a High-Molarity Tris Buffer System without Urea. Anal. Biochem. 155, 83–88 (1986).

59. Li, N. & Jarvis, R.P. Recruitment of Cdc48 to chloroplasts by a UBX-domain protein in chloroplast-associated protein degradation. Nat. Plants (2024).

60. Kikuchi, S. et al. Uncovering the Protein Translocon at the Chloroplast Inner Envelope Membrane. Science 339, 571–574 (2013).

61. Schindelin, J., et al. Fiji: an open-source platform for biological-image analysis. Nat. Methods 9, 676–682 (2012).

62. Nickel, C., Brylok, T. & Schwenkert, S. In Vivo Radiolabeling of Arabidopsis Chloroplast Proteins and Separation of Thylakoid Membrane Complexes by Blue Native PAGE. Methods Mol Biol 1450, 233–45 (2016).

63. Karimi, M., De Meyer, B. & Hilson, P. Modular cloning in plant cells. Trends Plant Sci. 10, 103–105 (2005).

64. Gasteiger, E. et al. ExPASy: The proteomics server for in-depth protein knowledge and analysis. Nucleic Acids Res. 31, 3784–8 (2003).

65. Wu, F.H. et al. Tape-Arabidopsis Sandwich - a simpler Arabidopsis protoplast isolation method. Plant Methods 5, 16 (2009).

66. Kovacheva, S., Bédard, J., Wardle, A., Patel, R. & Jarvis, P. Further studies on the role of the molecular chaperone, Hsp93, in plastid protein import. Plant J. 50, 364–379 (2007).

67. Punjani, A., Rubinstein, J.L., Fleet, D.J. & Brubaker, M.A. cryoSPARC: algorithms for rapid unsupervised cryo-EM structure determination. Nat. Methods 14, 290–296 (2017).

68. Rohou, A. & Grigorieff, N. CTFFIND4: Fast and accurate defocus estimation from electron micrographs. J. Struct. Biol. 192, 216–21 (2015).

69. Yang, B. et al. Identification of cross-linked peptides from complex samples. Nat. Methods 9, 904–906 (2012).

70. Lu, L. et al. Identification of MS-Cleavable and Noncleavable Chemically Cross-Linked Peptides with MetaMorpheus. J. Proteome Res. 17, 2370–2376 (2018).

71. Wassenaar, T.A., Ingólfsson, H.I., Böckmann, R.A., Tieleman, D.P. & Marrink, S.J. Computational Lipidomics with *insane*: A Versatile Tool for Generating Custom Membranes for Molecular Simulations. J. Chem. Theory Comput. 11, 2144–2155 (2015).

72. Berendsen, H.J.C., Postma, J.P.M., Vangunsteren, W.F., Dinola, A. & Haak, J.R. Molecular-Dynamics with Coupling to an External Bath. J. Chem. Phys. 81, 3684–3690 (1984).

73. Abraham, M.J. et al. GROMACS: High performance molecular simulations through multi-level parallelism from laptops to supercomputers. SoftwareX 1–2 19–25 (2015).

74. Humphrey, W., Dalke, A. & Schulten, K. VMD: Visual molecular dynamics. J Mol. Graph. 14, 33–38 (1996).

75. Gowers, R.J. et al. MDAnalysis: A Python package for the rapid analysis of molecular dynamics simulations. In S. Benthall and S. Rostrup, editors, Proceedings of the 15th Python in Science Conference, pages 98-105, Austin, TX. SciPy.

76. Hunter, J.D. Matplotlib: A 2D graphics environment. Comput. Sci. Eng. 9, 90–95 (2007).

77. Deutsch, E.W. et al. The ProteomeXchange consortium at 10 years: 2023 update. Nucleic Acids Res. 51, D1539–D1548 (2023).

