## Supplementary Information for "Assembly of two functionally-distinct protein import complexes in the outer membrane of plant chloroplasts"

#### SUPPLEMENTARY FIGURES

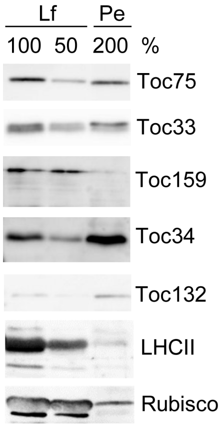


**Supplementary Fig. 1. Analysis of TOC complex components in petals.**

Total protein extract from the petals (Pe) of 6-week-old WT plants was analysed by immunoblotting alongside the chloroplast-enriched leaf fraction (Lf) described in **Fig. 1e**. Twice the amount (200%) of the petal sample was loaded relative to the chloroplast-enriched sample, so as to achieve normalized levels of the Toc75 component. Two different loading amounts (%) of the Lf sample are shown.

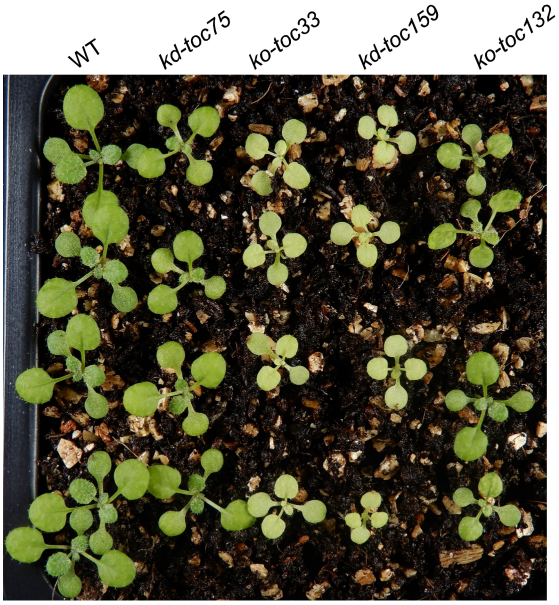


**Supplementary Fig. 2. Visible phenotypes of the *toc* mutants used in this study.**

The *kd-toc75*, *ko-toc33*, *kd-toc159* and *ko-toc132-2* mutants were grown alongside WT plants under identical conditions. The plants were photographed at the age of three weeks. Other names used previously to describe these mutants are as follows: *mar1/toc75-III-3* (*kd-toc75*), *ppi1-1* (*ko-toc33*), *fts1/ppi2-3* (*kd-toc159*), and *toc132-2* (*ko-toc132*).

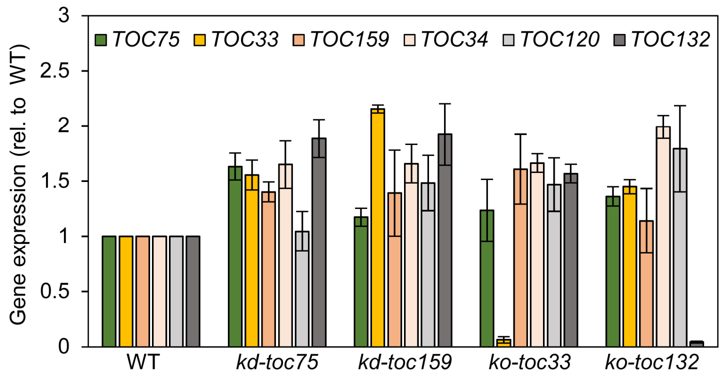


**Supplementary Fig. 3. Analysis of TOC component gene expression in the *toc* mutants.**

Gene expression analysis was performed by qRT-PCR using RNA samples extracted from 2-week-old plants of the indicated genotypes. Expression data for *TOC* genes were normalized using data for *ACTIN2*. All values are means ± SEM (n = 3 experiments). The data show that *TOC* genes (other than those directly affected by the mutation) are not generally under-expressed in *toc* mutants. The *TOC75* mutation in *kd-toc75* (causing missence mutation G658R) and the *TOC159* mutation in *kd-toc159* (causing premature stop at codon 1472) do not negatively affect transcription levels of *TOC75* and *TOC159*, respectively; though these mutations do strongly affect protein accumulation **(Fig. 2)**.

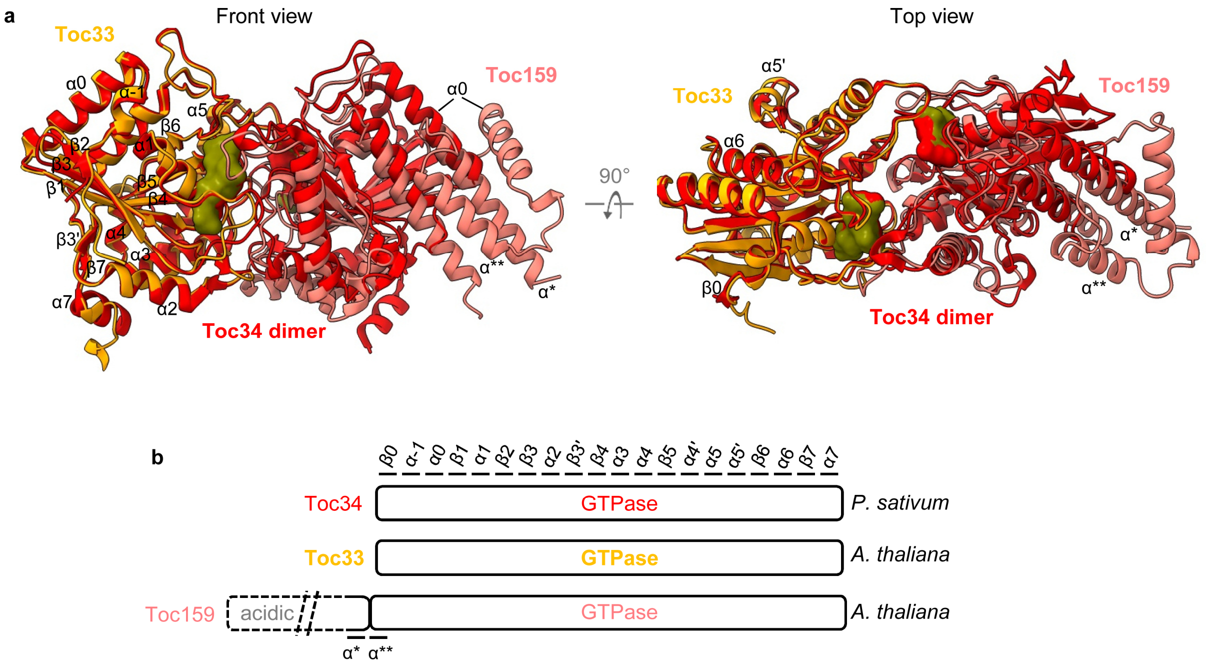


**Supplementary Fig. 4. Similarity of the Toc33-Toc159 GTPase heterodimer to the crystal structure of a pea Toc34 homodimer.**

**a.** Superposition of the GTPase heterodimer from the AF3-generated TOC-P structure with the published crystal structure of a *P. sativum* Toc34 homodimer (PDB: 1H65)[30]. *Arabidopsis* Toc33 and Toc159 are shown in orange and salmon pink, respectively; the pea Toc34 homodimer is shown in red; and the two GTPs (depicted as surface models) are shown in olive green. Left side, front view; right side, top view.

**b.** Secondary structural elements of the *P. sativum* Toc34 GTPase domain as previously described[30]. These structural elements were largely conserved in the GTPase domains of Toc33 and Toc159. Two upstream sequences of the Toc159 GTPase domain were additionally predicted by AF3 to be α-helices, as shown (α*, α**).

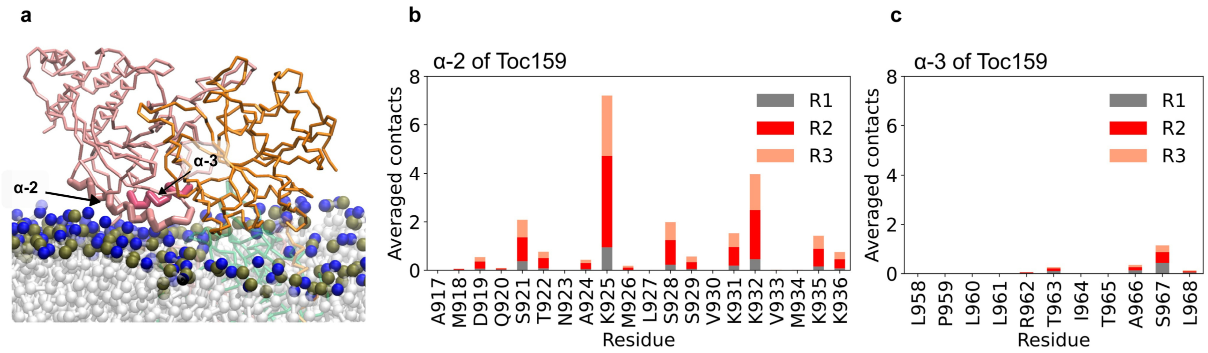


**Supplementary Fig. 5. Molecular dynamics simulations reveal the membrane contact sites of cytosolic regions of Toc159.**

**a.** Membrane-contacting cytosolic regions of Toc159 and Toc33 are shown. The helices α-2 (salmon pink) and α-3 (dark pink) of Toc159 are depicted with thick lines. POPC molecules are shown in light grey, with their headgroup moieties represented as coloured spheres (phosphate, khaki green; choline, blue).

**b,c.** Analysis of the lipid-α-2 and lipid-α-3 contacts. Bar charts showing averaged contacts between the α-2 or α-3 helices of Toc159 and POPC. The total contact counts for each residue along the whole production run were divided by the number of frames, producing an average value per frame. A cut-off of 0.6 nm was used to define a direct contact between any protein and POPC bead.

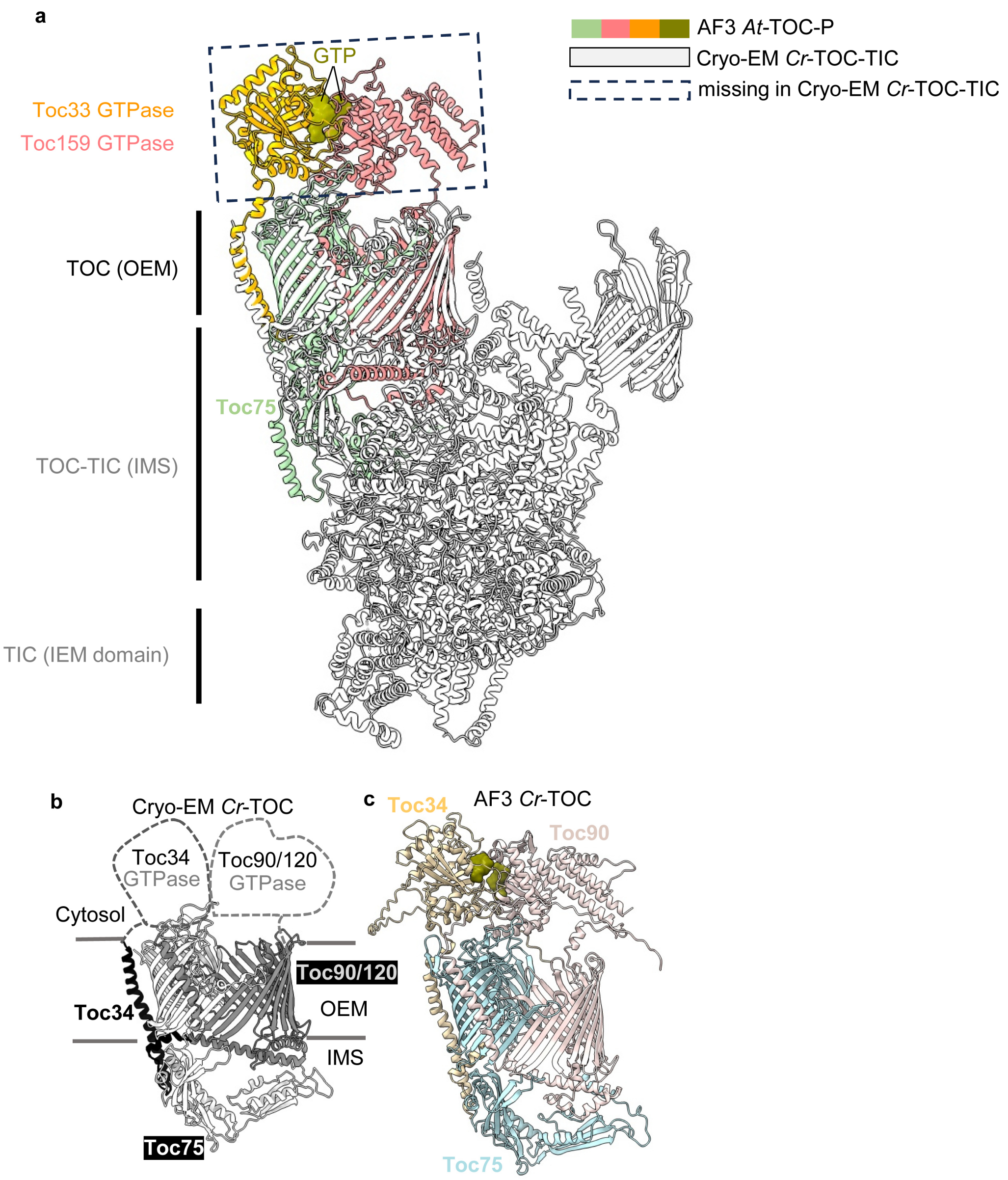


**Supplementary Fig. 6. Comparison of the Arabidopsis TOC-P AF3 structure with an algal TOC-TIC cryo-EM structure.**

**a.** Superposition (front view) of the AF3-generated TOC-P complex (coloured) with a cryo-EM structure of the *C. reinhardtii* TOC-TIC supercomplex (grey); just one of the published algal cryo-EM structures is used here (PDB: 7VCF) as these structures are very similar[27,28]. This analysis revealed similarities in the arrangement of the membrane-embedded β-barrel domains, but also some clear differences between the two structures. Most notably, while the GTPase domains of Toc33 and Toc159 were depicted with high confidence in the AF3 model of the *Arabidopsis* TOC-P complex, the GTPase domains of the algal receptors (termed Toc34 and Toc120/Toc90) were entirely missing, in both of the cryo-EM structures[27,28].

**b.** Cryo-EM structure of a TOC complex from *C. reinhardtii* (PDB: 7VCF). The absence of resolution regarding the cytosolic GTPase domains of the receptors (Toc34 and Toc90/120) is highlighted.

**c.** AF3-predicted structure of the same *C. reinhardtii* TOC complex revealing similarity with the corresponding cryo-EM structure (compare with **b**), and showing that the arrangement of the GTPase domains is similar to that in Arabidopsis TOC-P (compare with **a**).

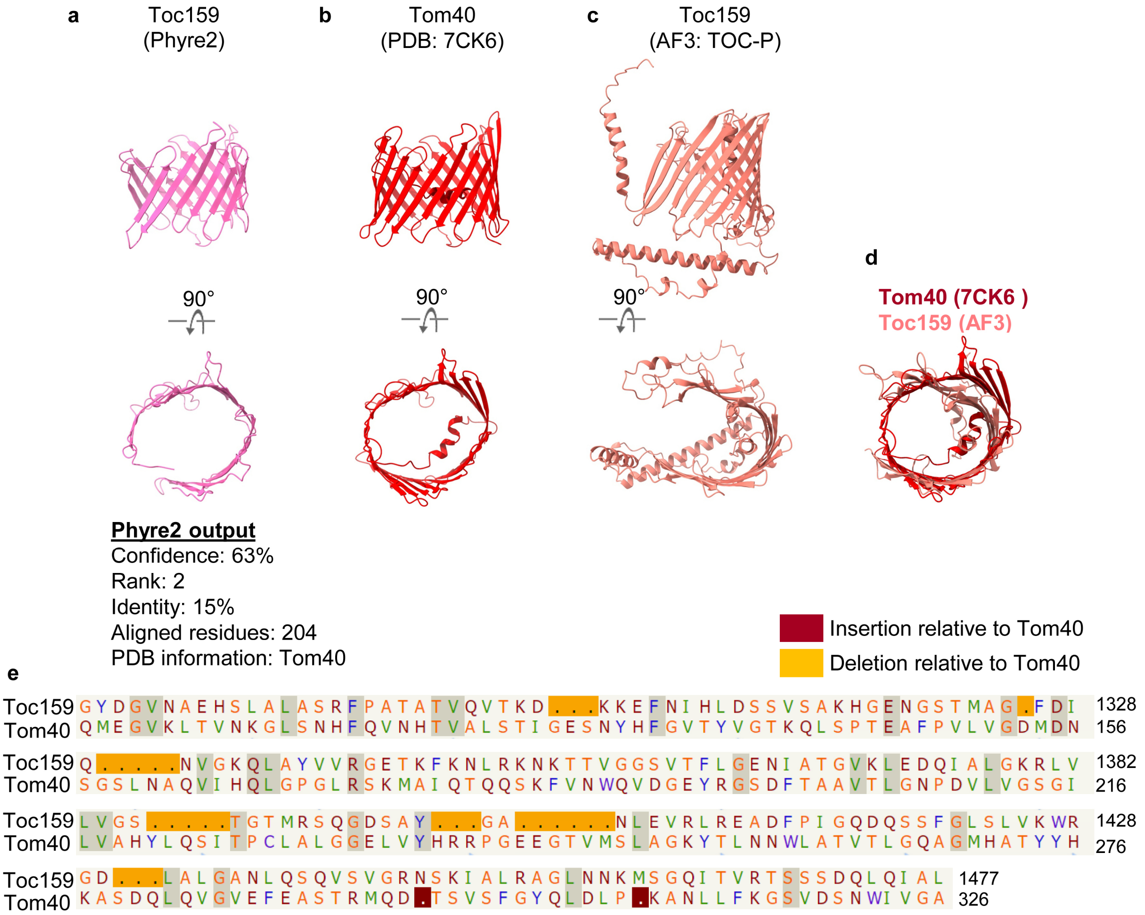


**Supplementary Fig. 7. Phyre2 identifies similarity between the Toc159 β-barrel and the mitochondrial protein import channel Tom40.**

The amino acid sequence of the membrane β-barrel domain of Toc159 (residues 1045-1503) was submitted to the Phyre2 protein threading server[52] to identify similar database structures. The second most similar hit identified (the first one being a homologous chloroplast translocon receptor from algae) was the mitochondrial protein import channel Tom40 (PDB: 7CK6). We compared the Toc159 structural model built by Phyre2 (using the Tom40 structure as a template; confidence, 63%) (**a**) with the original Tom40 structure (**b**), and with the membrane domain of Toc159 from our AF3-generated TOC-P structure **(see Fig. 3)** (**c**). In each case, front and top-down views are shown; outputted data from Phyre2 are also shown in **a**. Superposition (top view only) of the Tom40 structure with our AF3-generated Toc159 β-barrel structure supports the structural similarity identified by Phyre2 (**d**). The amino acid sequence alignment generated by Phyre2, covering the area of Toc159-Tom40 structural similarity, is also shown (**e**); identical residues are shaded. Together, these analyses revealed that the overall β-sheet folding pattern within the β-barrel domain of Toc159 (14 strands) is similar to that seen in the larger β-barrel of Tom40 (19 strands), in relation to both the arrangement of the β-strands and the size and arrangement of the inter-strand loops. It is noteworthy that both barrels (Toc159 and Tom40) have an open structure in the sense that the channel openings are not occluded.

**SUPPLEMENTARY DATA BELOW ARE APPENDED TO THIS FILE* OR PROVIDED AS SEPARATE FILES†:**

***Supplementary Fig. 8. Mass spectra identifying TOC crosslinks.**

The MS/MS spectra shown correspond to crosslinks between two peptides within Toc75 (#1-4) or Toc159 (#5-7). The crosslink spectra of #1, #2, #3 and #6 are from MetaMorpheus 1.0.9, and those of #4, #5 and #7 are from pLINK3.

**†Supplementary Video 1. Front view from molecular dynamics simulation of the TOC-P complex.**

TOC-P dynamics during a full 10 μs long simulation (run R3). Protein backbones are shown as sticks, coloured by chain following the same colour scheme as in **Fig. 3c** (Toc33, orange; Toc159, salmon pink; Toc75, green). Linker regions are highlighted in light blue. The membrane and solvent are omitted for clarity.

**†Supplementary Video 2. Top-down view from molecular dynamics simulation of the TOC-P complex.**

TOC-P dynamics during a full 10 μs long simulation (run R3). Protein backbones are shown as sticks, coloured by chain following the same colour scheme as in **Fig. 3c** (Toc33, orange; Toc159, salmon pink; Toc75, green). Linker regions are highlighted in light blue. The membrane and solvent are omitted for clarity.

**†Supplementary Table 1. List of peptides including BS3 crosslinks identified by LC-MS/MS.**

***Supplementary Table 2. Primers used in this study.**

**a.** Primers used to generate the *HA-Toc75* transgene.
**b.** Primers used for *TOC* gene expression analysis by RT-PCR.
**c.** Primers used to construct transit peptide fusions of *SSU* or *E1α* to *YFP/CFP* for protoplast expression.
**d.** Primers used for gene expression analysis by RT-PCR following protoplast expression.

#### Supplementary Fig. 8 (part i)

**Toc75 (Crosslink #1)**

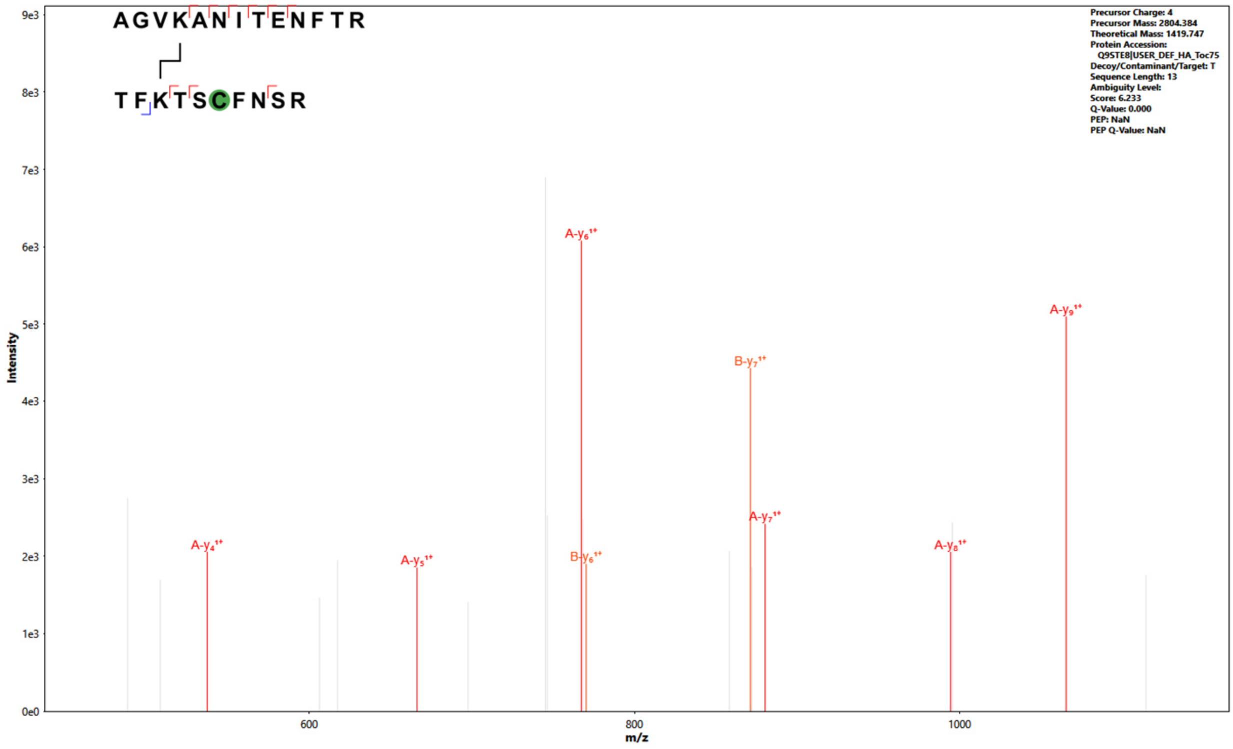


**Toc75 (Crosslink #2)**

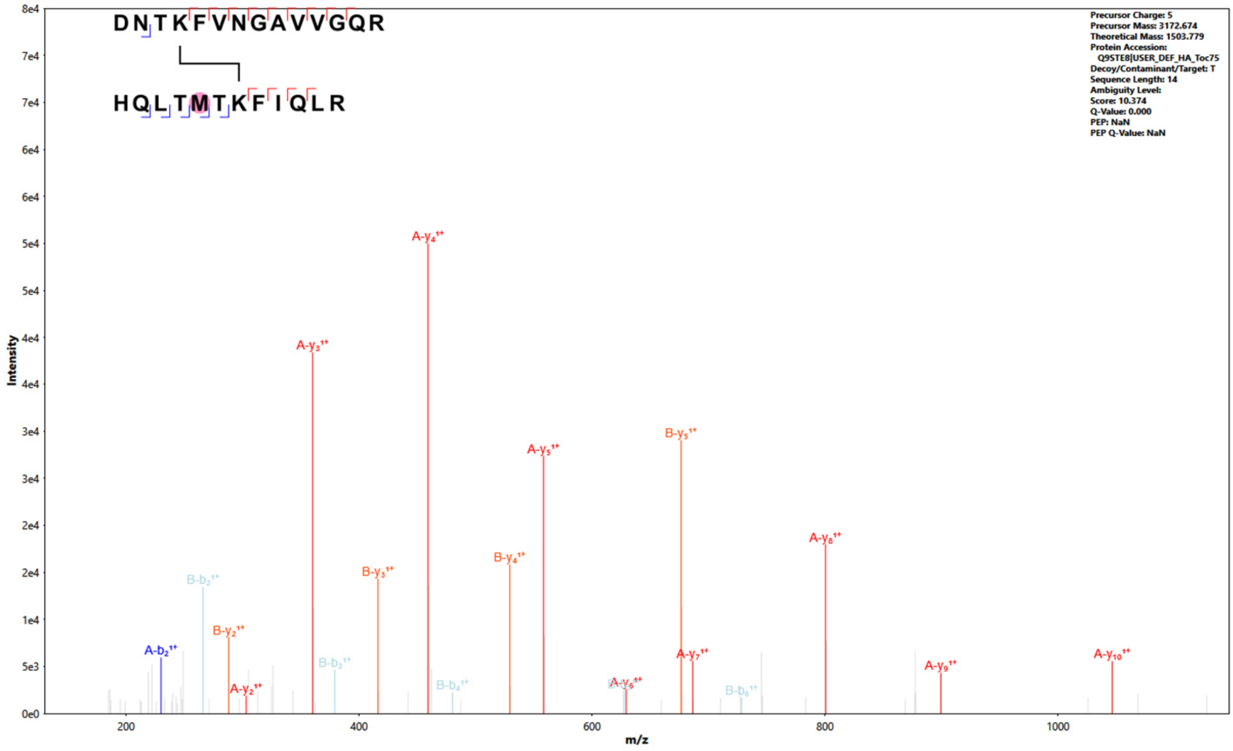

#### Supplementary Fig. 8 (part ii)

##### Toc75 (Crosslink #3)

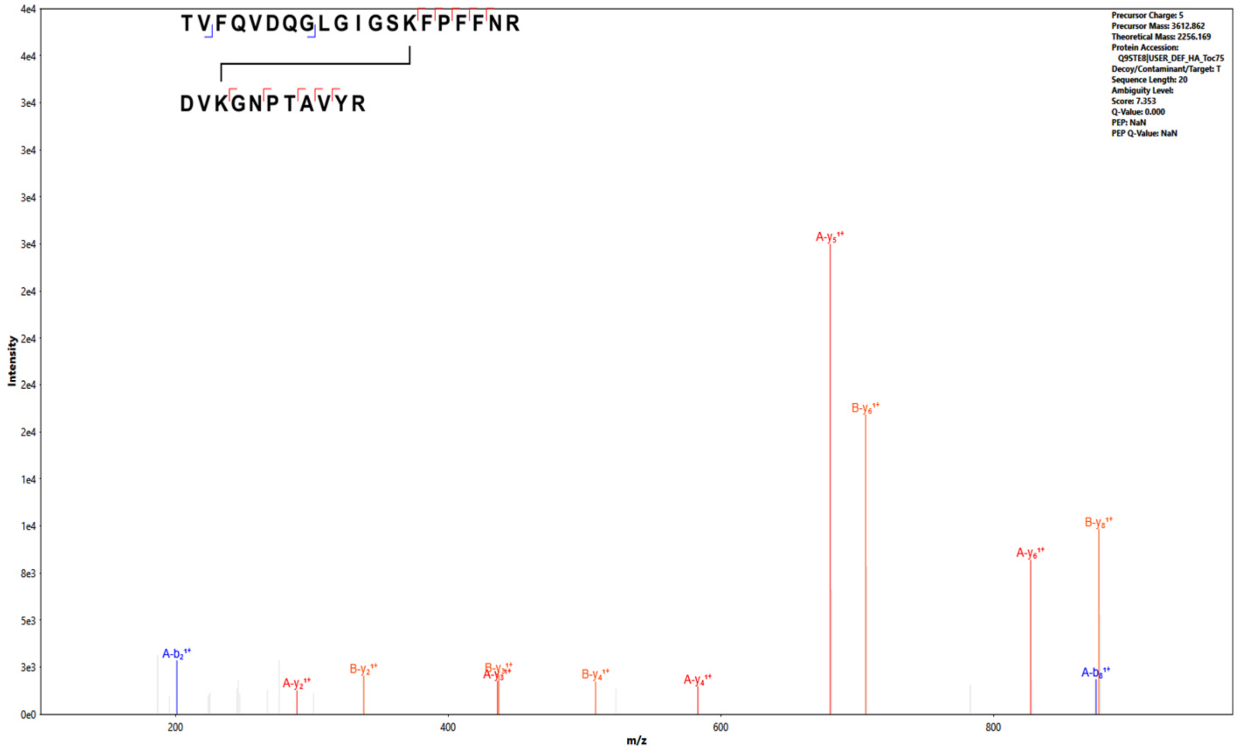


##### Toc75 (Crosslink #4)

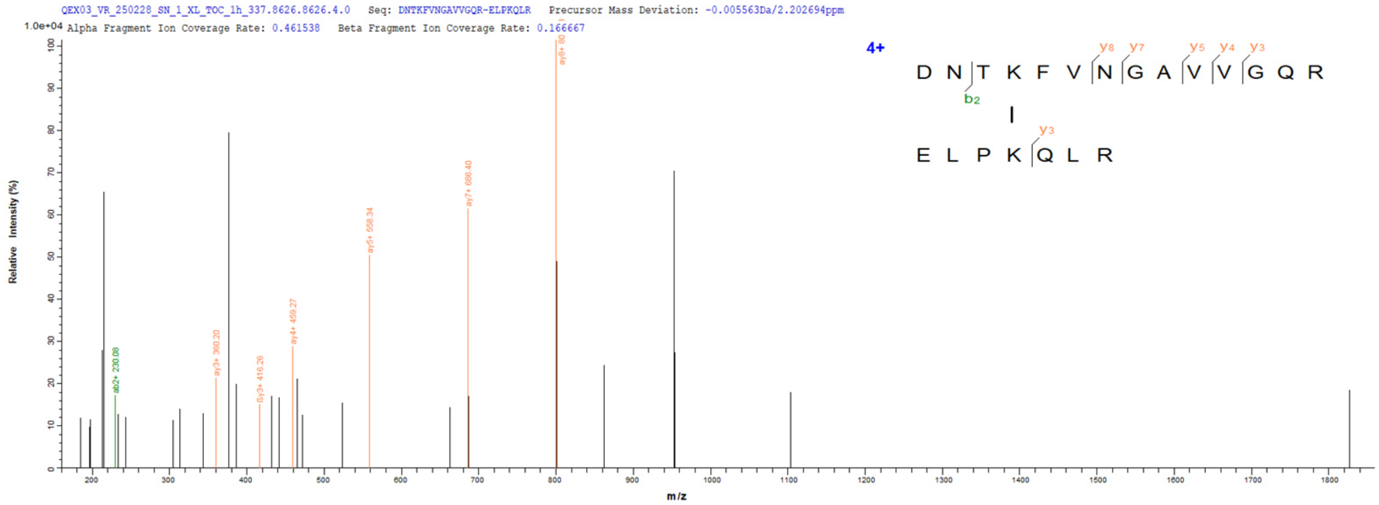


##### Toc159 (Crosslink #5)

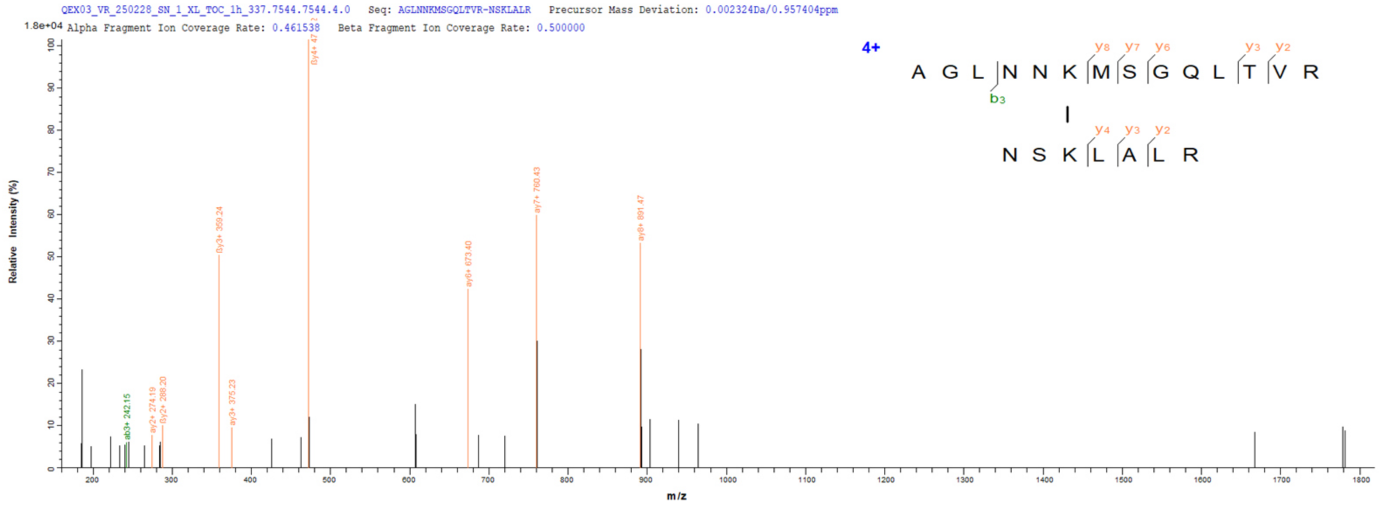

#### Supplementary Fig. 8 (part iii)

**Toc159 (Crosslink #6)**

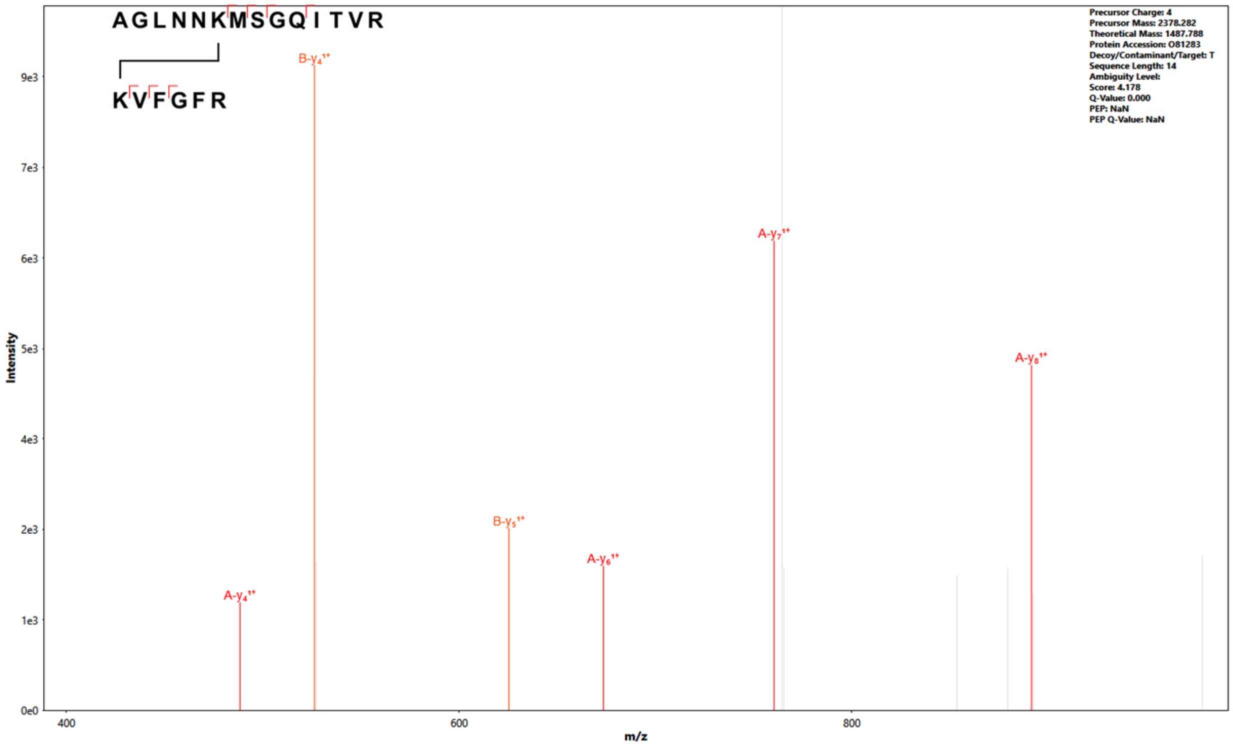


**Toc159 (Crosslink #7)**

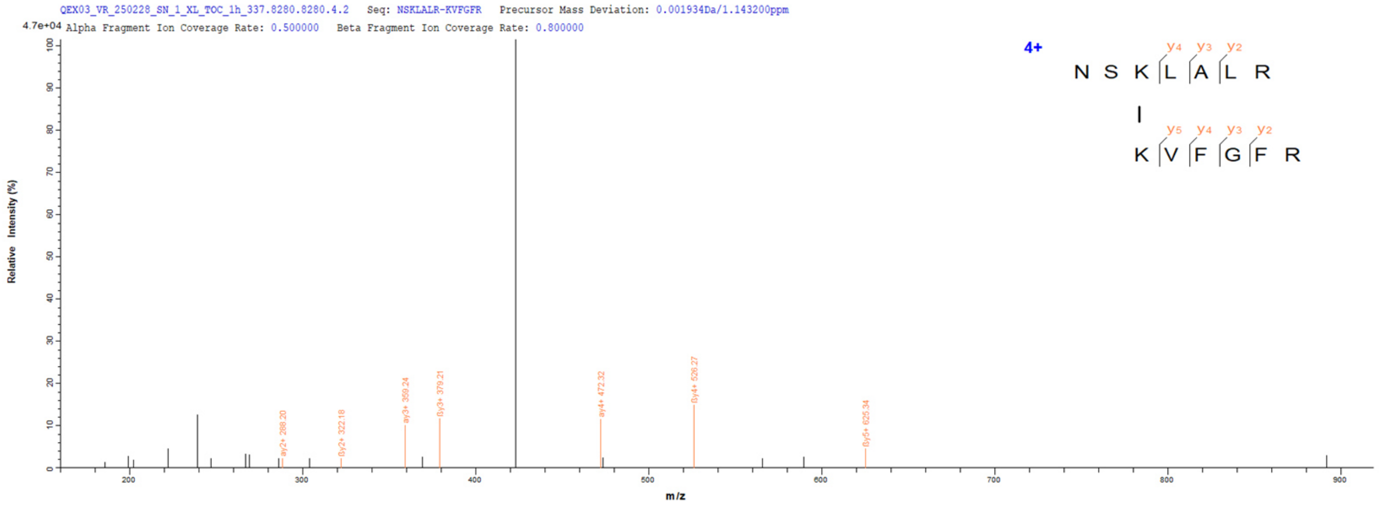

### Supplementary Table 2. Primers used in this study.

#### a. Primers used to generate the *HA-Toc75* transgene.

| Primer name | Primer sequence (5' to 3') | Used to generate… | Figure(s) |
|---|---|---|---|
| gToc75-F | CATGTCTCTTCTTGGTGCTGAGA | *TOC75* genomic fragment | Fig. 1a |
| gToc75-R | CAGCATCTACCATCTCAACAGAATC | | |
| mToc75-HA-F | TTCCTGATTATGCTGAACAATCACCGGA | *HA-tag* coding sequence | |
| mToc75-HA-R | CATCATAAGGATATTCATCAGCAACCGC | | |

#### b. Primers used for *TOC* gene expression analysis by RT-PCR.

| Primer name | Primer sequence (5' to 3') | Used to analyse… | Figure(s) |
|---|---|---|---|
| Toc75-F | AGTCAATCCACGCCCAGATG | *TOC75* | Fig. 4a,<br>Extended Data Fig. 9,<br>Supplementary Fig. 3 |
| Toc75-R | GGCTGGAATGAAGCCAATGTG | | |
| Toc159-F | CAACCAACCCCTTCTACGCT | *TOC159* | |
| Toc159-R | CCTTGGCACTACTCACAGCA | | |
| Lhcb1-F | TCGAGTGAGAGACAGGAGGA | *LHCB1* | |
| Lhcb1-R | ACAACGGAGTGAACCCAAGA | | |
| Toc33-F | ACAAGTCGTCCGTGTCAGTC | *TOC33* | |
| Toc33-R | CACAAGTCCAGGGGTGTCAA | | |
| Toc34-F | AGGTGGTGTTGGAAAGTCATCA | *TOC34* | |
| Toc34-R | AGACCAAGGTAGGCCTCAGT | | |
| Toc120-F | ACCGATGTGTTGCAAGAGGA | *TOC120* | |
| Toc120-R | TGGCTCTCTCCAATCTCCGA | | |
| Toc132-F | TTGAAGAGGCAGTGGGTGAT | *TOC120* | |
| Toc132-R | AAATTCAGCCTCTCCCGCAC | | |
| Actin2-F | TCAGATGCCCAGAAGTCTTGTTCC | *ACTIN2* | |
| Actin2-R | CCGTACAGATCCTTCCTGATATCC | | |

#### c. Primers used to construct transit peptide fusions of *SSU* or *E1α* to *YFP/CFP* for protoplast expression.

| Primer name | Primer sequence (5' to 3') | Used to generate… | Figure(s) |
|---|---|---|---|
| TP-RbcS1A-F (SSU) | AAAAAGCAGGCTCCATGGCTTCCTCTATGCTC | *TP-SSU* coding sequence | Fig. 7a,b, Extended Data Fig. 10a,b |
| TP-RbcS1A-R (SSU) | AGAAAGCTGGGTTAGGAAGGTAAGAGAGAGTC | | |
| TP-E1α-F | AAAAAGCAGGCTCCATGGCGACGGCTTTCGCTCC | *TP-E1α* coding sequence | |
| TP-E1α-R | AGAAAGCTGGGTTCAGGCTGGTATTATTGGTGG | | |

#### d. Primers used for gene expression analysis by RT-PCR following protoplast expression.

| Primer name | Primer sequence (5' to 3') | Used to analyse… | Figure(s) |
|---|---|---|---|
| eIF4E1-F | AAACAATGGCGGTAGAAGACACTC | *eIF4E1* | Extended Data Fig. 10c,d |
| eIF4E1-R | AAGATTTGAGAGGTTTCAAGCGGTGTAAG | | |
| SSU-F | CAATAATACCAGCCTGAACCCA | *SSU-YFP/CFP* *E1α-YFP/CFP* | |
| E1α-F | TCTCTCTTACCTTCCTAACCCA | | |
| YFP/CFP-R | TTACTTGTACAGCTCGTCCATG | | |
